# Auditory brainstem models: adapting cochlear nuclei improve spatial encoding by the medial superior olive in reverberation

**DOI:** 10.1101/694356

**Authors:** Andrew Brughera, Jason Mikiel-Hunter, Mathias Dietz, David McAlpine

## Abstract

Listeners perceive sound-energy as originating from the direction of its source, even as direct sound is followed milliseconds later by reflected sound from multiple different directions. Early-arriving sound is emphasised in the ascending auditory pathway, including the medial superior olive (MSO) where binaural neurons encode the interaural time difference (ITD) cue for spatial location. Behaviourally, weighting of ITD conveyed during rising sound-energy is stronger at 600 Hz, a frequency with higher reverberant energy, than at 200 Hz where reverberant energy is lower. Here we computationally explore the combined effectiveness of adaptation before ITD-encoding, and excitatory binaural coincidence detection within MSO neurons, in emphasising ITD conveyed in early-arriving sound. With excitatory inputs from adapting model spherical bushy cells (SBCs) of the bilateral cochlear nuclei, a Hodgkin-Huxley-type model MSO neuron reproduces the frequency-dependent emphasis of rising vs. peak sound-energy in ITD-encoding. Maintaining the adaptation in model SBCs, and adjusting membrane speed in model MSO neurons, hemispheric populations of model SBCs and MSO neurons, with simplified membranes for computational efficiency, also reproduce the stronger weighting of ITD information conveyed during rising sound-energy at 600 Hz compared to 200 Hz. This hemispheric model further demonstrates a link between strong weighting of spatial information during rising sound-energy, and correct unambiguous lateralisation of reverberant speech.

## Introduction

Sound propagating directly from its source to a listener’s ears is typically followed milliseconds later by multiple reverberant copies arriving from different directions (Fig. 1). Despite this mixture of direct and reflected sound pressure, generating non-stationary spatial information in the binaural cues—interaural time differences (ITDs) and interaural intensity differences (IIDs)—listeners typically perceive a sound as punctate, and originating from the direction of its source (Dietz et al., 2013). Perception of the true location of the source can persist even when the intensity of reflected sound matches or exceeds that of early-arriving, direct sound (Haas, 1951), facilitating ‘cocktail party listening’: attending to a single talker against a background of many competing voices (Cherry, 1953).

**Fig. 1.**
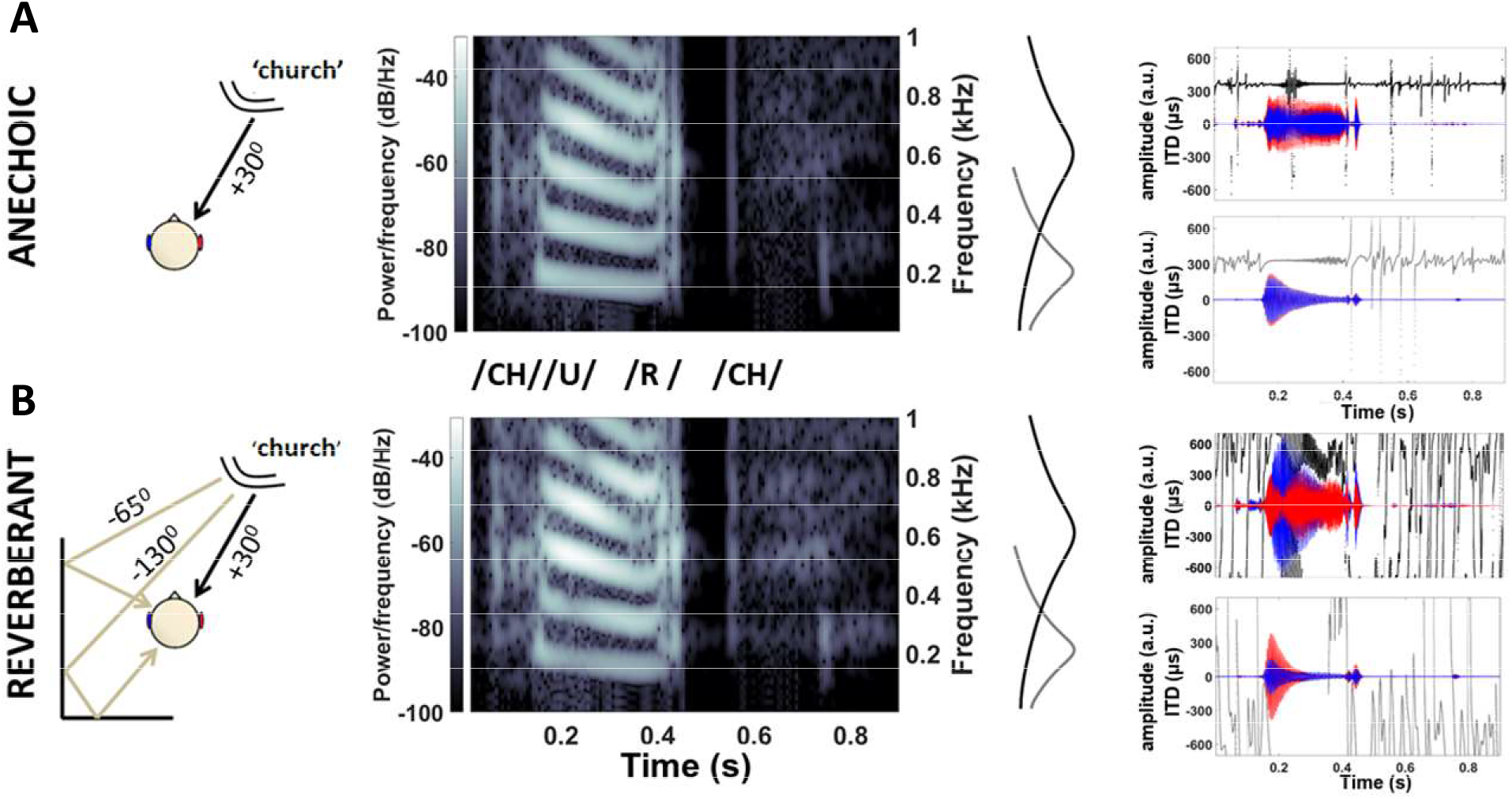
Reverberant copies of direct speech produce confounding binaural cues. **A** Anechoic speech stimulus ‘church’ (direct sound, without reverberation), spoken at 70 dB SPL by a female talker located +30° to the right and front of a virtual listener. Spectrogram shows speech energy <1 kHz at left ear. Gammatone filters centred at 200 Hz (light grey) and 600 Hz (dark grey) show effects of cochlear filtering on speech waveforms in the left (blue) and right (red) ears. Instantaneous ITDs (light/dark grey) (restricted to ±700us, human physiological range) were consistently near +363 μs. **B** Reverberant speech stimulus including direct speech plus two simulated reflections from listener’s left, the first from −65° delayed by 4 ms, and the second from −130° delayed by 8 ms. Reverberant copies increased the energy at the left ear (note the generally brighter spectrogram above 400 Hz). Reverberant energy extends into the quiet pause between the vowel and final consonant (0.45-0.55 ms), generating rapidly varying instantaneous ITDs (also restricted to ±700us), along with conflicting ILD cues in the 600-Hz channel

Behavioural emphasis of early-arriving ITD information is frequency dependent (Hu et al., 2017): for amplitude-modulated 600-Hz sounds, listeners are more sensitive to ITDs conveyed during the rising-energy portion than they are to ITDs at the energy peak; this is not the case at 200 Hz, where listeners are equally sensitive to ITDs conveyed during rising and peak energy. Acoustically, reverberant energy can be 20 dB less intense at 200 Hz than at 600 Hz in many outdoor settings (Traer & McDermott, 2016). For both frequencies, listeners are least sensitive to ITD during falling sound-energy, when direct sound is most likely to be corrupted by reverberation. These data suggest that spatial auditory brain mechanisms transmit reliable information, and suppress unreliable information, accounting for natural frequency-profiles in reverberant energy which decrease below 1500 Hz.

Neural emphasis of early-arriving sound is observed in the auditory nerve, brainstem, midbrain, and auditory cortex (Dietz et al., 2014; Fitzpatrick et al., 1999; Liebenthal & Pratt, 2002; Litovsky & Yin, 1998a, 1998b). In the brainstem, medial superior olive (MSO) neurons, performing binaural coincidence detection that encodes ITDs in low-frequency sounds (Mathews et al., 2010; Yin & Chan, 1990), respond strongest to their preferred ITD during rising sound-energy in amplitude-modulated binaural beats (AMBBs), producing an emphasis of early-arriving spatial information that is consistent with adaptation in monaural projection pathways to binaural MSO neurons (Dietz et al., 2014). Within these pathways, MSO neurons receive bilateral excitation from spherical bushy cells (SBCs) of the ventral cochlear nuclei (VCN) (Smith et al., 1993). Each SBC is driven by 1-3 auditory nerve fibres (ANFs), each terminating in a calyceal Endbulb of Held synapse (Lorente De No, 1981). Potential adaptive mechanisms include: spike-rate adaptation in ANFs (Moser & Beutner, 2000; Zilany & Carney, 2010); short-term plasticity (STP, synaptic depression) observed *in vitro* at the synapse from ANF to SBC (Oleskevich et al., 2000; Wang & Manis, 2008; Yang & Xu-Friedman, 2009, 2015); and glycinergic inhibition at SBCs observed *in vivo* (Keine & Rübsamen, 2015; Keine et al., 2016; Kuenzel et al., 2011, 2015).

Here we present simplified computational models of the auditory brainstem (Figs. 2 and 5), exploring the combined effectiveness of monaural adaptation, and excitatory binaural coincidence detection, in emphasising ITD conveyed in early-arriving sound, and improving the lateralisation of speech in reverberation. In our simplified models, adapting ANFs (Zilany et al., 2014, 2009) drive SBCs (Rothman & Manis, 2003c) that adapt according to the STP *in vitro*; despite weak STP *in vivo* (Keine et al., 2016; Kuenzel et al., 2011), nearly identical temporal properties to the inhibition *in vivo* (Kuenzel et al., 2015; Wang & Manis, 2008) support expected adaptive effects. With excitatory inputs from the adapting model SBCs, a Hodgkin-Huxley-type model MSO neuron reproduces *in vivo* AMBB-responses of MSO neurons and the frequency-dependent emphasis of rising vs. peak sound-energy in ITD-encoding (Dietz et al., 2014; Hu et al., 2017). Maintaining the adaptation in model SBCs, and adjusting membrane speed in model MSO neurons within the observed range (Bondy & Golding, 2018; Remme et al., 2014; Scott et al., 2007), hemispheric populations of model SBCs and MSO neurons, with simplified membranes for computational efficiency, also reproduce the stronger weighting of ITD conveyed during rising sound-energy at 600 Hz compared to 200 Hz. This hemispheric model further demonstrates a link between strong weighting of spatial information during rising sound-energy, and correct unambiguous lateralisation of reverberant speech (Haas, 1951).

**Fig. 2.**
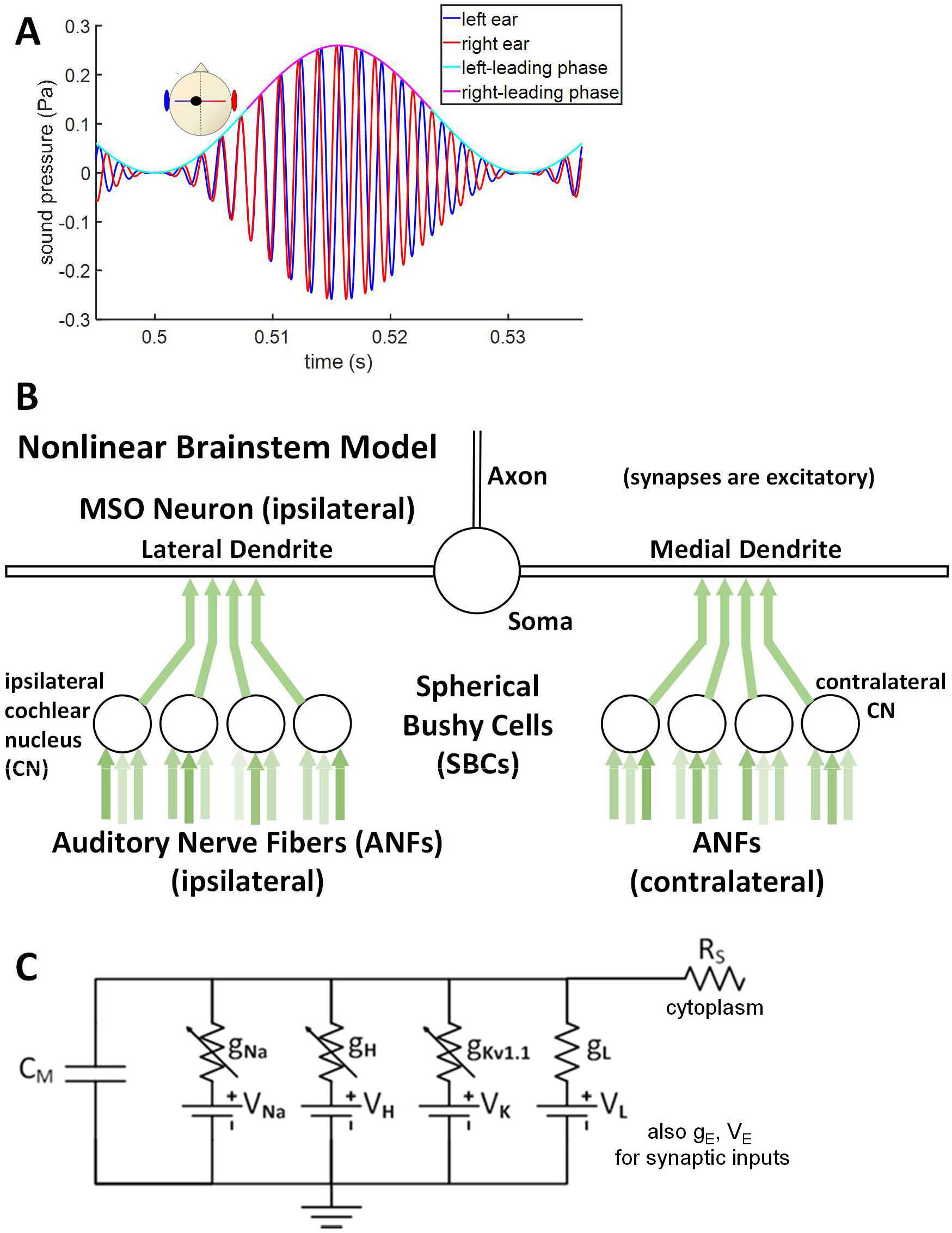
Amplitude-modulated binaural beats (AMBBs) and the nonlinear brainstem model **A** An AMBB stimulus: the AM rate is set equal to binaural-beat frequency (the difference in frequency between right and left ears). Presented via headphones, interaural phase difference (IPD) cycles through 360° at the same rate as AM. *Start-IPD* (the IPD at zero amplitude) is a free parameter. Shown with right-carrier 616 Hz, left-carrier 584 Hz, AM 32 Hz, and *start-IPD* 270° (right channel trailing by 90°), resulting in zero IPD at the midpoint of rising-amplitude. **B** Nonlinear brainstem model for an MSO neuron and its excitatory inputs, with adaptive spike-failures in SBCs phenomena-logically modelled using synaptic depression. **C** Sub-compartment in the nonlinear brainstem model (see Methods)

**Fig. 5.**
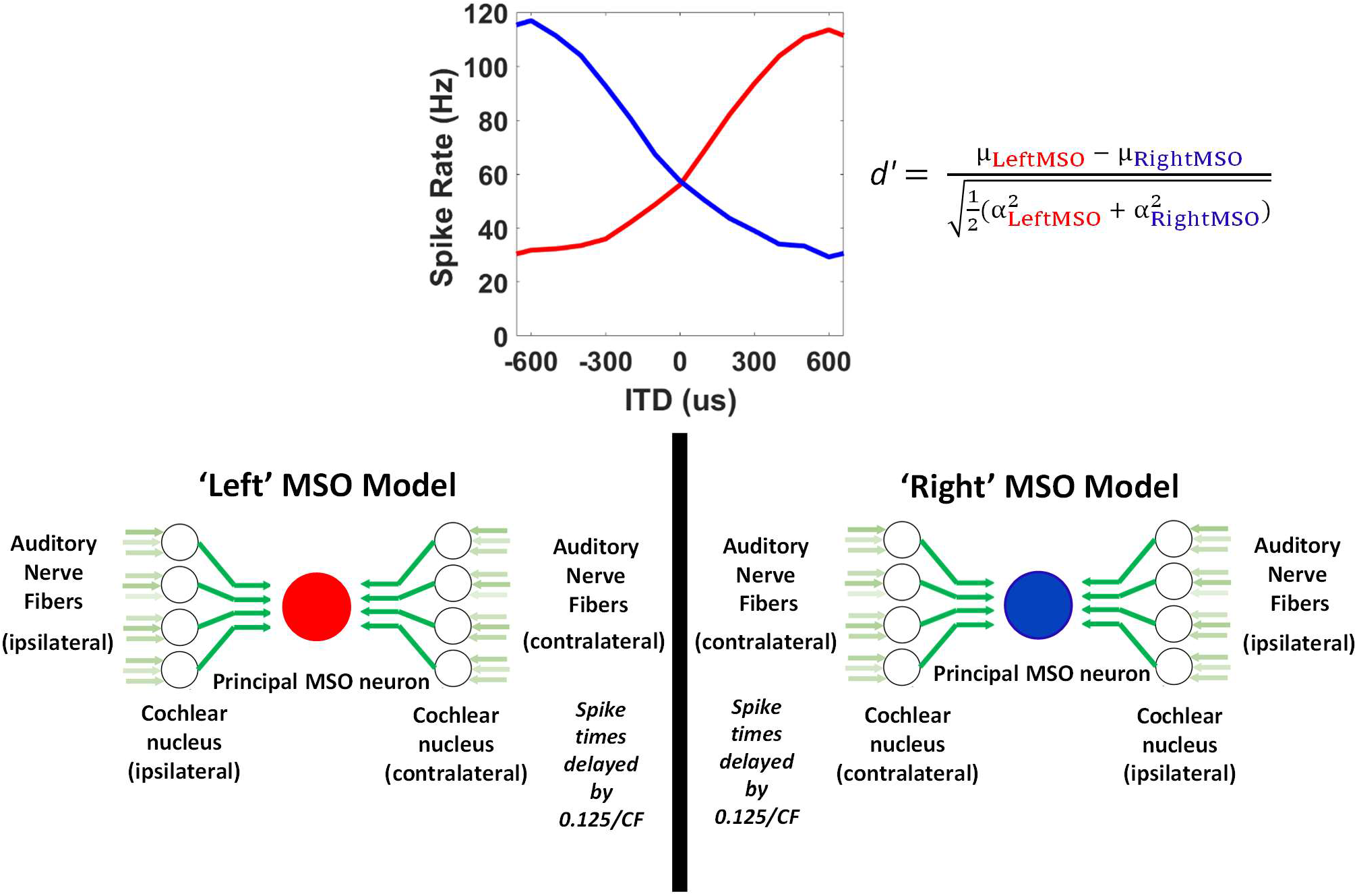
Overview of the hemispheric-difference model consisting of two MSO populations (red—left MSO and blue—right MSO), each containing fifty linear, single-compartment model neurons, with either relatively fast or slow membranes (Remme et al., 2014). Each MSO neuron model receives four excitatory inputs ‘bilaterally’ from linear, single-compartment model SBCs of the CN. Three independently-simulated, individually-depressing and medium-spontaneous-rate model ANFs (Zilany et al., 2014) provide excitatory drive to each SBC model neuron

## Methods

### Nonlinear brainstem model

This computational model (Fig. 2B) incorporates an MSO neuron and its excitatory input pathways, beginning with an auditory periphery model for humans (Glasberg & Moore, 1990; Zilany et al., 2014), with 12 left and 12 right adapting model ANFs of medium spontaneous rate. Three model ANFs drive each adapting model SBC of the VCN (Rothman & Manis, 2003c; Rudnicki & Hemmert, 2017). Four left and four right SBCs project excitatory synaptic inputs to a multi-compartment model MSO neuron (Brughera et al., 2013). Acoustic stimuli and the auditory periphery model are implemented in Matlab (Natick, Massachusetts, USA) (www.mathworks.com). Model SBCs and MSO neuron are Hodgkin-Huxley-type (Hodgkin & Huxley, 1952), implemented in the Python 3.7 Brian2 Neural Simulator (Stimberg et al., 2019). Code is available at Github: https://github.com/AndrewBrughera/Mso_SbcStp_EE1 Data, analysis scripts, and code are available at figshare: https://doi.org/10.6084/m9.figshare.11955219.v1

#### Acoustic Stimuli

Acoustic stimuli are amplitude-modulated binaural beats (AMBBs) (Fig. 2A) (Dietz et al., 2013) at 75 dB SPL RMS at peak amplitude, in which binaurally presented tones of a slightly different frequency are modulated in amplitude at a rate equal to the difference in frequency across the two ears. With this stimulus, each cycle of rising and falling sound-energy contains a full cycle of interaural phase disparities.

#### Auditory Periphery Model

The acoustic stimuli are processed by an auditory periphery model for humans (Glasberg & Moore, 1990; Zilany et al., 2014). Peripheral processing includes 24 adapting ANFs of medium spontaneous rate: 12 each in the left and right ears, with a 200-Hz or 600-Hz characteristic frequency (CF, the frequency at which a neuron fires above spontaneous rate for the lowest sound pressure level, SPL). CF results from the frequency-tuning of the inner ear, and the CFs of ANFs distally driving a neuron.

#### Model SBCs

SBCs (Type II VCN neurons) are modelled as Hodgkin-Huxley-type point neurons (Rothman & Manis, 2003c), adjusted for temperature 37°C) with three independent, excitatory synapses each driven by a model ANF (Lorente De No, 1981). Each model synapse is an excitatory synaptic conductance 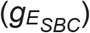 in series with an excitatory reversal potential of 0 mV. Each input spike causes a variable increment in its synapse’s excitatory conductance (with a standard unadapted maximum increment, or maximum excitatory synaptic strength, 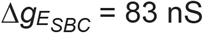; spike-threshold was 34 nS for a single input from rest, using the same model membrane and faster synapses at 38°C (Rothman & Manis, 2003c)). 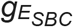 then decays exponentially with time-constant 0.2 ms (Kuenzel et al., 2011, 2015). The increment (i.e., the synaptic strength) is variable due to depression at the individual synapses, modelled as in Rudnicki & Hemmert (2017): immediately after an input spike and its associated increment in excitatory conductance, the synaptic strength is multiplied by 1 – *u,* where *u* = 0.5. This synaptic depression recovers exponentially with time-constant 25 ms, as measured *in vitro* (Wang & Manis, 2008). Low synchrony to amplitude modulation (AM) in the model SBCs is consistent with the auditory nerve at high SPL (Joris & Yin, 1992) and lower synchrony in primary-like neurons of the cochlear nuclei (Rhode & Greenberg, 1994). Compared with a single input, three inputs per model SBC increased spike rates, reduced synchrony to AM, and supported a small number of excitatory inputs (Couchman et al., 2010) to the model MSO neuron.

#### Model MSO neuron

The Hodgkin-Huxley-type model principal MSO neuron has separate compartments representing the bilateral dendrites, soma, and axon (Zhou et al., 2005). The model axon functions simply as a spike generator, without myelination or Nodes of Ranvier. Compared with a previous model (Brughera et al., 2013), the soma is simplified being spherical and iso-potential, and the somatic and dendritic membrane conductances for voltage-sensitive ion-channels are scaled by 0.6. Models for fast-acting ion-channels, the low-threshold potassium (K_LT_) channels (Mathews et al., 2010) and sodium (Na) channels (Scott et al., 2010) are based on the MSO. The model for slowly-varying hyperpolarization-activated cyclic nucleotide (H) channels remains based on the VCN (Rothman & Manis, 2003c).

On the bilateral model dendrites, eight excitatory synapses (Couchman et al., 2010) are located one each at 42.5, 47.5, 52.5, and 57.5% of the dendritic length. Each synapse is modelled as a variable conductance 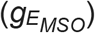 in series with an excitatory reversal potential (*V*_*E*_) of 0 mV. Noting that STP in the brainstem is weak *in vivo* (Keine et al., 2016; Kuenzel et al., 2011; Lorteije et al., 2009; Yin & Chan, 1990), the synapses are non-depressing. Each synapse is driven by a single model SBC from the same side. An input spike increments its synaptic conductance by a fixed amount (the excitatory synaptic strength, 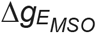), and the conductance then decays exponentially with time-constant *τ*_*E*_ = 0.4 ms, within physiological range (Fischl et al., 2012; Franken et al., 2015). With inputs from adapting SBCs (results in Figs. 3B and 4), 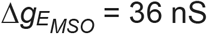, comparable with excitatory fibre conductances of 37 ± 4 nS measured *in vitro* (Couchman et al., 2010).

**Fig. 3.**
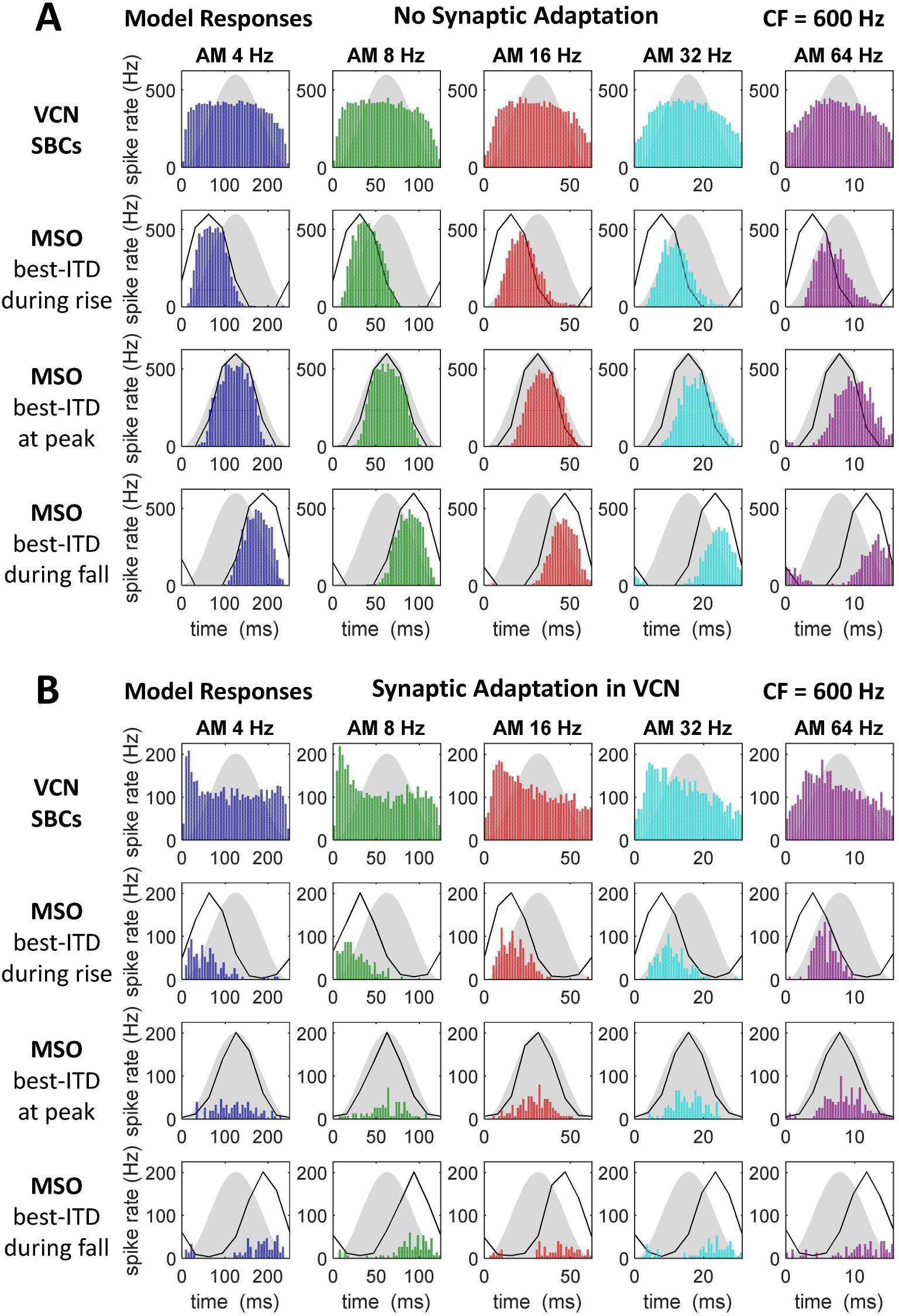
AMBB cycle histograms for nonlinear model brainstem neurons acoustically stimulated by sound-frequencies centred at CF 600 Hz: **A** Without synaptic adaptation, auditory nerve to SBC synapses were strongly supra-threshold. Model SBCs adapted slightly. Model MSO neuron responded to best ITD strongly at peak sound-energy, and slightly but consistently less strongly during rising and falling energy (3-way comparisons, *P* < 10^−9^). **B** With synaptic adaptation in the cochlear nuclei, the auditory nerve to SBC synapses were supra-threshold at full strength. Model SBCs adapted. Model MSO neuron responded to best ITD strongly during rising energy; weakly to moderately at peak; and weakly during falling, energy (3-way comparisons, *P* ≤ 0.003). *Definitions:* Grey silhouettes show the AM envelope. Black lines show static IPD functions. Best-ITD = 0. Best-ITD during rise means *start-IPD* = 270°. Best-ITD at peak means *start-IPD* = 180°. Best-ITD during fall means *start-IPD* = 90°

**Fig. 4.**
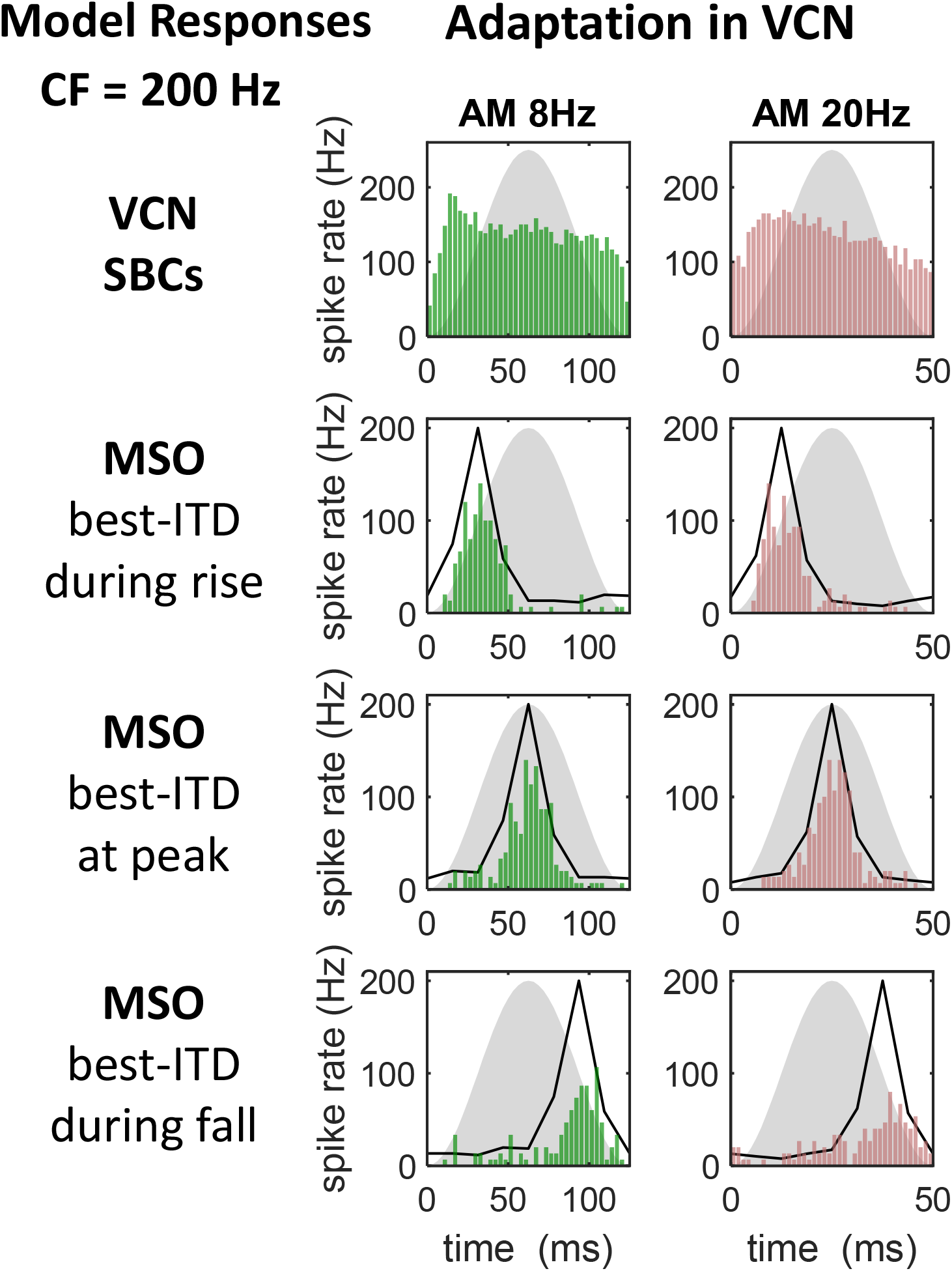
AMBB cycle histograms for nonlinear model brainstem neurons stimulated by frequencies centred at CF 200 Hz. Same synaptic adaptation and model SBCs as in Fig. 3B, adapted less at 200 Hz. Same fast model MSO neuron as in Fig. 3B: spike counts were not significantly different for best-ITD during rising vs. peak energy, and were higher for best-ITD at peak energy vs. falling energy (*P* < 0.01). *Definitions:* Grey silhouettes show the AM envelope. Black lines show static IPD functions. Best-ITD = 0. Best-ITD during rise means *start-IPD* = 270°. Best-ITD at peak means *start-IPD* = 180°. Best-ITD during fall means *start-IPD* = 90°

Membrane parameters for each compartment (Table 1) are given below. Certain parameters (Table 2) were adjusted for the conditions yielding results shown in Figs. 3-4. *V*_*AP-THRESHOLD*_ is a fixed threshold for counting action potentials: when the membrane voltage near the midpoint of the model axon transitions from −35 mV or less to greater than −20 mV, a spike is counted; this threshold is simply for counting, and does not affect the operation of the model. The somatic membrane time-constant of 0.39 ms was calculated from measured membrane impedance in the model (see below and Supplemental Fig. S1).

**Table 1.** Model MSO neuron: compartmental and membrane parameters.

| Parameter | Dendrites (2) | Soma | Axon |
| --- | --- | --- | --- |
| Temperature ($^{\circ}\text{C}$) | 37 | 37 | 37 |
| Number of sub-compartments | 20 | 2 | 51 |
| Diameter ($\mu\text{m}$) | 3.5 | 30 | 2 |
| Length ($\mu\text{m}$) | 150 | 30 | 400 |
| Resistivity, $R_S$ (ohm x cm) | 150 | 0 | 150 |
| $C_M$ ($\mu\text{F}/\text{cm}^2$) | 1 | 1 | 1 |
| $V_{KLT}$ (mV) | -106 | -106 | -106 |
| $V_{Na}$ (mV) | n/a | +62.1 | +62.1 |
| $V_H$ (mV) | -43 | -43 | -43 |
| $V_L$ (mV) | -65 | -65 | -65 |
| $g_{maxKLT}$ ($\text{S}/\text{cm}^2$) | 0.00132 | 0.0324 | 0.0595 |
| $g_{maxNa}$ ($\text{S}/\text{cm}^2$) | 0 (none) | 0.0432 | 0.25 |
| $g_{maxH}$ ($\text{S}/\text{cm}^2$) | 0.00066 | 0.01296 | 0.0025 |
| $g_L$ ($\text{S}/\text{cm}^2$) | 0.00005 | 0.00005 | 0.00005 |
| $V_E$ (mV) | 0 | n/a | n/a |
| $V_{AP-THRESHOLD}$ (mV, set) | n/a | n/a | -20 |
| $V_{REST}$ (mV, measured) | -60.53 | -60.52 | -64.35 |

**Table 2.** Excitatory synaptic parameters.

| Results shown in: | Fig. 3A | Figs. 3B & 4 |
| --- | --- | --- |
| Model SBCs | control | test |
| Full Synaptic Strength, $\Delta g_{ESBC}$ (nS) | 400 | 83 |
| Synaptic Time-Constant, $\tau_{ESBC}$ (ms) | 0.2 | 0.2 |
| Synaptic Depression, $u$ | 0 (none) | 0.5 |
| Recovery Time-Constant (ms) | n/a | 25 |
| Model MSO neurons | control | test |
| Synaptic Strength, $\Delta g_{EMSO}$ (nS) | 21 | 36 |
| Synaptic Time-Constant, $\tau_{EMSO}$ (ms) | 0.4 | 0.4 |

Within this model MSO neuron, its functional electric-circuit unit, the sub-compartment (Fig. 2C), connects to its neighbouring sub-compartments via series resistivity (R_S_) representing the neuron’s internal cytoplasm. Within each sub-compartment, the neural membrane is modelled as a transmembrane capacitance (*C*_*M*_) in parallel with ion-channel populations: Na, H, K_LT_, and leakage (L). Each ion-channel population is represented as a reversal potential (*V*_*Na*_, *V*_*H*_, *V*_*KLT*_, *V*_*L*_) in series with a conductance (*g*_*Na*_, *g*_*H*_, *g*_*KLT*_, *g*_*L*_). With the exception of a fixed leakage conductance (*g*_*L*_) representing voltage-insensitive ion channels, each conductance value is equal to a maximum conductance multiplied by voltage-sensitive activation and inactivation gating variables with integer exponents (Hodgkin & Huxley, 1952).

Kirchhoff’s current equation, applied to the model MSO neuron, states that the sum of currents entering any point is zero:

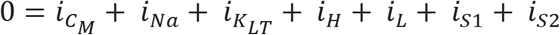

Series currents, *i*_*S*1_ and *i*_*S*2_, are calculated by the Brian2 simulator according to the voltage differences, resistivity, and geometry of the related sub-compartments.

Capacitive membrane current 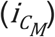 increases with membrane capacitance and the time-derivative of membrane potential *V*:

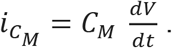

Leakage current: *i*_*L*_ = *g*_*L*_(*V* − *V*_*L*_).

Na current (Scott et al., 2010), which is rapidly varying, is based on the MSO:

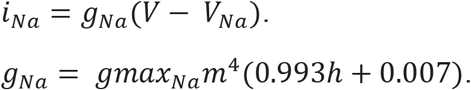

Each voltage-sensitive ionic conductance has a constant maximum conductance value (*gmax*) (Table 1), and has voltage and time dependencies defined by a subset of the activation and inactivation variables *m*, *h*, *w*, *z*, and *r*, each with a rate of change governed by a first-order differential equation with a time-constant divided by the *Q*_10_ temperature factor of 3^(*T* – 22)/10^, where *T* is set equal to human body temperature, 37°C. For Na channels:

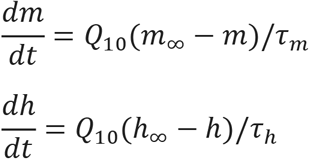

For model Na channels, steady-state activation (*m*_∞_) and inactivation (*h*_∞_), and their respective time constants (*τ*_*m*_ and *τ*_*h*_) in milliseconds, are functions of membrane potential in millivolts (Scott et al., 2010):

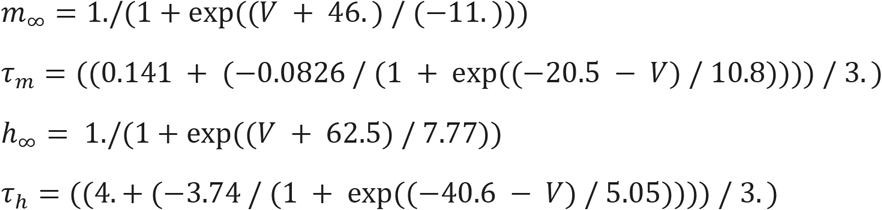

K_LT_ current (Mathews et al., 2010), which is rapidly-varying, is based on the MSO:

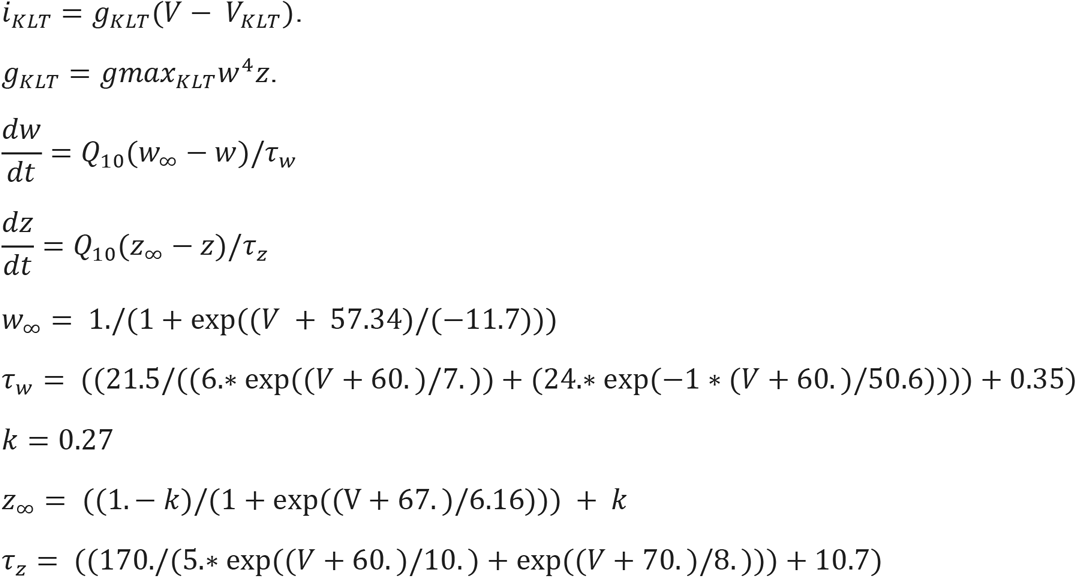

The non-inactivating H current (Rothman & Manis, 2003c), which is slowly-varying, remains based on the VCN:

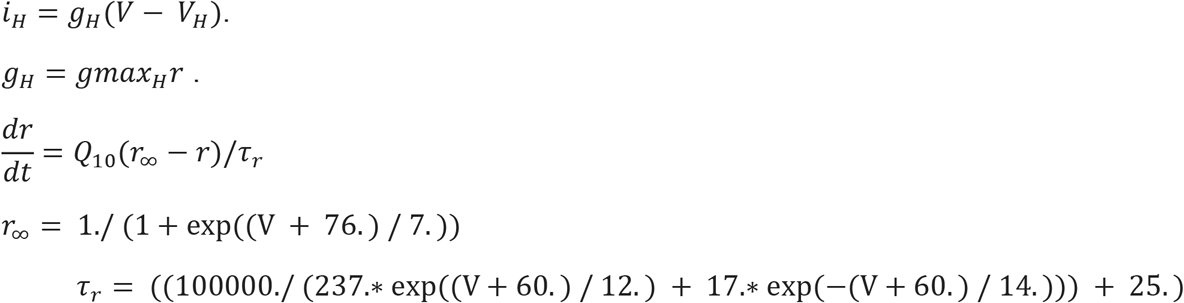

Maximum conductance (*gmax*) values are set for plausible resting potentials and membrane time constants. Leakage conductance equals the value of Scott et al. based on measurements from the MSO (Scott et al., 2010). The reversal potential for leakage in the model dendrites and soma is reduced from −60 mV to −65 mV, now consistent with the model axon (Brughera et al., 2013). For voltage-sensitive ion-channels, ratios of the maximum conductances for ion channels are calculated for resting potentials: near −60 mV in the soma and dendrites, consistent with physiological values promoting activation of model K_LT_ channels (Mathews et al., 2010); and near −64 mV in the axon to reduce inactivation of model Na channels. The 4-mV difference produces a small current from soma to axon, calculated at 20 picoamperes (pA), much less than synaptic currents. Although less negative than the −68 mV in a dedicated MSO axon model (Lehnert et al., 2014), the resting potential in our model axon has been shown to support large axonal action potentials, without significant back-propagation to the soma at its higher resting potential and lower ratio of Na to K_LT_ conductance (Brughera et al., 2013), which is consistent with the small somatic action potentials of MSO neurons (Scott et al., 2007). From the calculated ratios of maximum conductance, values are scaled. Model somatic membrane impedance as a function of frequency was measured by simultaneous injection of a transmembrane bias current, and a sinusoidal current containing a linear frequency sweep from 1 to 2000 Hz during a duration of 1 second (Hutcheon & Yarom, 2000; Puil et al., 1986; Remme et al., 2014). Prior to stimulus, the model membrane settled for 0.1 s to its resting potential, −60.52 mV. Settling was followed by a steady inward bias current applied for 0.9 s. The bias current was then maintained as the frequency sweep in membrane current (250-pA peak amplitude) was also applied. A resonance in membrane impedance emergences with increases in bias current (Supplemental Fig. S1). Assuming that the resonant membrane is a second-order system (Nilsson & Riedel, 2008), the membrane time-constant is equal to the reciprocal of angular resonance frequency. With a bias current of 300 pA, resulting in a holding potential of −59.22 mV, the resonance at 408 Hz indicates a membrane time-constant of 1/(2π × 408 Hz) = 0.390 ms, which is within the range of 0.3 to 0.6 ms measured in principal MSO neurons (Couchman et al., 2010; Scott et al., 2007).

#### AMBB period histograms

For each AMBB period histogram from the nonlinear model, spikes were counted in forty non-overlapping bins, each covering 1/40 of the AM cycle (unsmoothed). Spike rates were calculated by dividing the spike count by the total time duration for each bin, across the multiple periods of eight different stimulus presentations of 0.75-second duration.

#### Eight starting-phases at decrements of 45° efficiently implemented IPDs

At each carrier and modulation-frequency combination, 8 acoustic stimuli, each with 1 of 8 carrier *starting-phases* (0°, −45°, −90°, …, −315°), were applied at each model ear, driving ANFs. For computational efficiency, *start-IPD* (the interaural phase difference at zero amplitude) was achieved by pairing spike times from each *starting-phase* in one ear, with the spike times from the other ear having the appropriate difference in *starting-phase*. While maintaining the proper *start-IPD*, each AMBB period histogram pooled spikes resulting from stimuli using the 8 *starting-phases* spanning each carrier period. Thus whilst the histograms show the effects of *start-IPD*, they do not show phase-locking to fine structure of the carriers.

In this study, “best-ITD during rise” denotes *start-IPD* = 270°; “best-ITD at peak” denotes *start-IPD* = 180°; and “best-ITD during fall” denotes *start-IPD* = 90°.

#### Chi-squared tests for significant differences in model spike counts

For each condition and modulation frequency, a chi-squared test for one and two degrees of freedom (2-way and 3-way comparisons) compared spike counts from the nonlinear model MSO neuron stimulated with AMBBs with best-ITD occurring during rising vs. peak vs. falling amplitude. Each chi-squared test yielded a probability (*P*) of the null hypothesis that the differences in spike count occurred randomly.

#### Synchrony index and Rayleigh statistic for significant phase-locking

For each modulation frequency and *starting-phase* combination, the synchrony index (*SI, R*) of spike times (*t*_*i*_) (Johnson, 1980) with respect to their phase *θ*_*i*_ within the AM cycle (with modulation frequency *f*_*m*_, and modulation period *T*_*m*_ = 1/*f*_*m*_) was calculated:

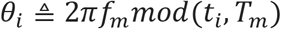

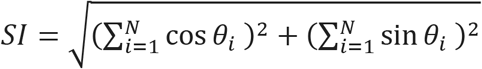, where N is the spike count.

The Rayleigh significance statistic (*2NR*^*2*^) was then calculated and converted to a *P* value, the probability of the null hypothesis that the AM period-histogram of spike times resulted from a uniform distribution (Rhode, 1976):

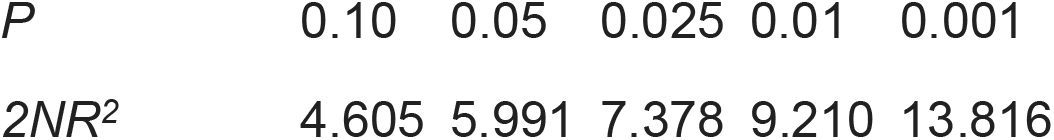

### Lateralisation model with Linear-membrane models

To study how monaural adaptation and biophysical profiles could impact lateralisation of acoustic stimuli in anechoic and reverberant conditions, we constructed a model of lateralisation whose brainstem stages (SBCs and MSO neurons) are represented by point neurons with linear biophysical properties (Remme et al., 2014). Data, analysis scripts, and code are available at figshare: https://doi.org/10.6084/m9.figshare.9899018.v1

Two stimuli were presented to the model: the first, a SAM tone with non-zero ITDs inserted at different phases of the AM cycle, as used behaviourally to determine how carrier frequency affects ITD sensitivity over the time course of an AM cycle (Hu et al., 2017); the second, a natural speech stimulus with early reflections, has been proposed as a simple simulation of a reverberant environment where correct lateralisation is only possible if ITD cues in the onset waveform are processed preferentially (Dietz et al., 2014).

#### SAM tones with non-zero ITDs inserted at different phases of AM cycle

This stimulus was generated according to the Hu et al. (2017) study that first used it. In summary, SAM tones were created from pulse trains that had been bandpass-filtered and then amplitude-modulated. Non-stationary ITDs were added by temporally shifting individual pulses in one of either the rising, peak or falling phases of the AM cycle. For the purposes of this study, non-stationary ITDs of +300 μs and +150 μs were inserted into SAM tones of either 200-Hz or 600-Hz carrier frequency respectively. These ITD values were chosen based on individual behavioural thresholds in the Hu et al. (2017) study. At each carrier frequency, four conditions were tested: two different AM frequencies (8 Hz or 20 Hz) and two different proportions of rising, peak or falling AM phases with a non-zero ITD inserted (20% or 40%). These parameters match those tested behaviourally by Hu et al. (2017).

#### A natural speech stimulus with early reflections

A single monosyllabic consonant-vowel nucleus-consonant (CNC) word token, ‘church’, (spoken by an Australian-English, female voice), was selected as a stimulus (Fig. 1 and Fig. 8). To simulate its arrival from a location one metre away to the right of the midline in an anechoic environment, the word’s waveform (normalised at 70dB) was convolved with a large pinnae HRTF for +30° azimuthal location (CIPIC database, UC Davis). A reverberant space was simulated by adding to the direct (+30°) pathway two early reflections off virtual walls stood behind and to the right of the subject (Fig. 1). These reflected copies of the words arrived from −65° and −130° angles (again convolved with large-pinnae HRTFs from the CIPIC database) and were delayed by 4ms and 8ms respectively to reproduce their elongated paths (Dietz et al., 2014).

**Fig. 8.**
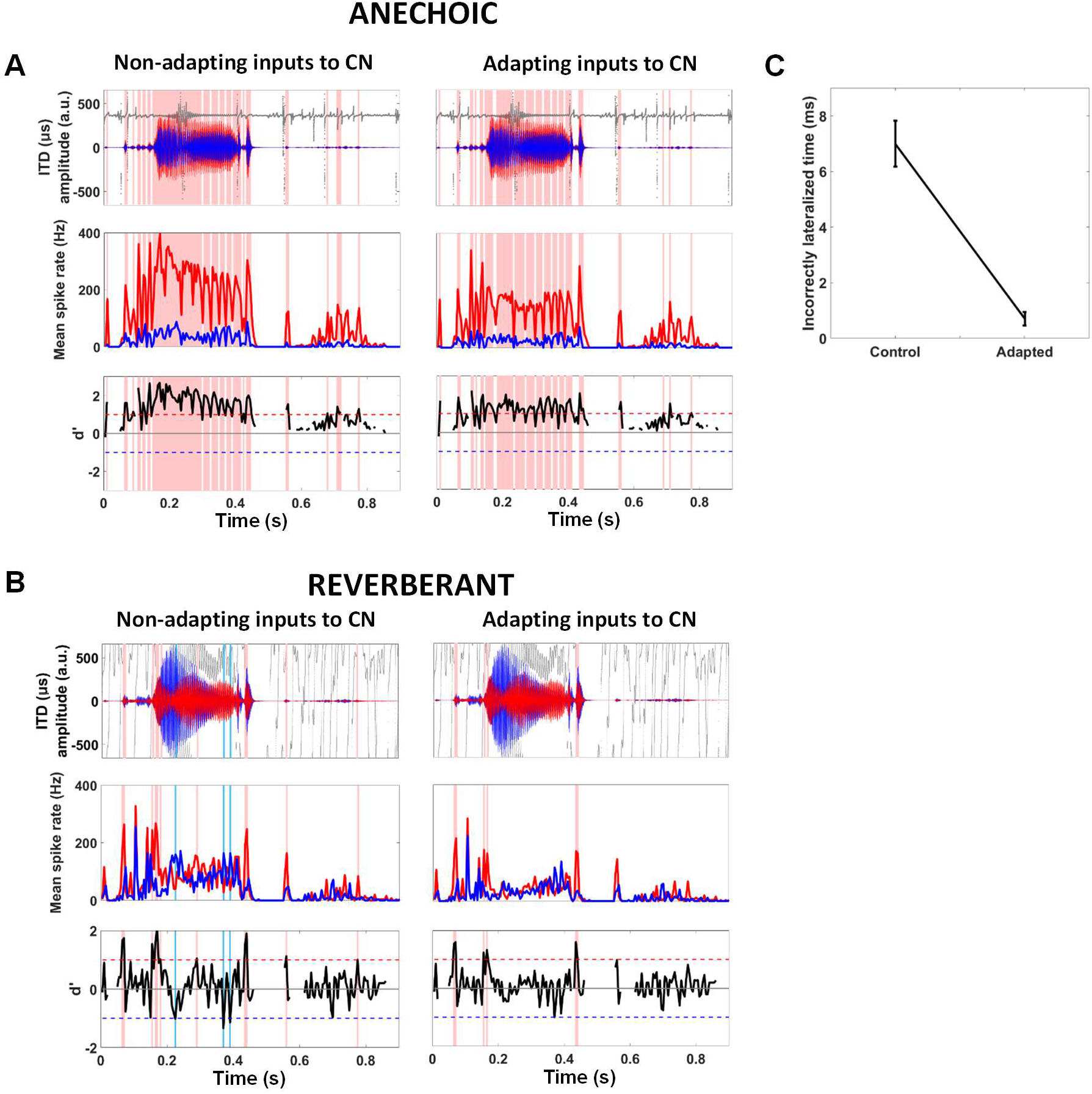
Lateralisation of ‘anechoic’ and ‘reverberant’ speech stimuli by the hemispheric-difference model (fast MSO, 600-Hz channel) with and without synaptically adapting inputs from the auditory nerve to the CN: **A** Anechoic stimulus is correctly lateralised to the right (pink vertical bands) for long periods independently of non-adapting (left column) or adapting (right column) synaptic inputs. Dry speech stimulus gammatone-filtered at 600 Hz with instantaneous ITDs (top row), mean firing rates of left (middle row, red) and right (middle row, blue) MSOs and d’ neuro-metric (bottom row) are displayed. **B** Reverberant stimulus requires adapting synaptic inputs (right column) for correct lateralisations alone (pink vertical bands). Non-adapting synaptic inputs to the cochlear nuclei produced both correct (left column, pink vertical bands) and incorrect lateralisations (left column, light blue vertical bands) due to reverberant stimulus’ highly variable ITDs and confounding ILDs (top row). The inclusion of synaptic adaptation from the auditory nerve to the cochlear nuclei (using the same auditory nerve simulations) removes all incorrect lateralisations. Remaining correct lateralisations correspond with stimulus onsets (right column, pink vertical bands). **C** Quantifying incorrect lateralisations with non-adapting and adapting synaptic inputs (same auditory nerve simulations for both) over 100 presentations of the reverberant speech stimulus. Synaptic adaptation produces a five-fold decrease in incorrect lateralisations (N = 100, *t*(99) = 9.51, *P* = 1.31 × 10^−15^, two-tailed paired T-test)

#### Linear model circuitry

The linear model, consists of three stages: an initial auditory periphery stage (the middle/inner ear and ANFs) and two brainstem stages (cochlear nuclei and MSO). Unlike the nonlinear model, the linear model incorporates two parallel circuits, representing the ‘left’ and ‘right’ MSO nuclei and their inputs. The difference in output rate between these parallel circuits is utilised to calculate lateralisation of the natural stimulus. The only discernible difference between ‘left’ and ‘right’ circuits is the contralateral 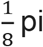 shifts in their MSO neurons’ IPD tuning curve peaks which are generated by delaying the outputs from the auditory periphery stage ipsilateral to the MSO nucleus in question. This shift is considered physiological given experimentally observed binaural tuning curves from the auditory brainstem whose optimal characteristics have been corroborated theoretically (McAlpine et al., 2001).

#### Auditory periphery stage

Peripheral processing of the acoustic stimuli in the linear model is performed by a cat middle/inner ear model (Zilany et al., 2014) in Matlab. Outputs from the auditory periphery are restricted to 200-Hz/600-Hz for the pulse-shifted SAM tones (matching their SAM tone’s carrier frequency) and 600-Hz for lateralisation of the speech stimulus with early reflections. This latter value was chosen based on it being within the 500-750 Hz frequency range in which temporal-fine-structure (TFS) ITD sensitivity is considered strongest (Ihlefeld & Shinn-Cunningham, 2011). When exploring effects, of synaptic depression at the inputs from auditory nerve to VCN, on the lateralisation of a speech stimulus, the same auditory nerve simulations are used for both the depressing and non-depressing inputs.

#### Linear-membrane model properties

Single-compartment, linear-membrane models are used to represent both SBC and MSO neurons in the linear model. These linear-membrane models can incorporate two dynamic currents (Remme et al., 2014): I_*W*_, a resonant current and/or, I_*n*_, an amplifying current. The current balance equation for the general neural model is as follows:

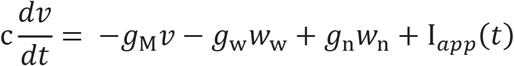

Where *c* is capacitance (picofarads), *g*_*M*_ is total membrane conductance (nanosiemens, nS) and I_app_ is the current (picoamperes) applied to the neural model. The resonant and amplifying currents, I_*W*_ and I_*n*_, are described by their conductances, *g*_*W*_ and *g*_*n*_ (in nS); the dynamics of their gating variables, *W*_*W*_ and *W*_*n*_ (in millivolts) are described generically by the equation:

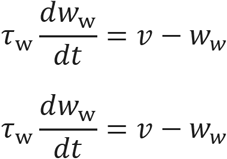

Where *W*_*x*_ is the gating variable whose associated time-constant is τ_*x*_ (milliseconds).

Two parameter sets are chosen to represent the range of membrane speeds recently observed in MSO neurons (Bondy & Golding, 2018; Remme et al., 2014). These example model membranes can be broadly characterised as either fast with a high resonance-frequency or slow with a low resonance-frequency as previously modelled by Remme et al. (2014). While both fast and slow model membranes are used to represent MSO neurons, only the fast model membrane was chosen to represent all SBCs,

#### Synaptic properties and plasticity

Providing excitatory drive to each linear model SBC, again 3 independently-simulated ANFs were applied. Excitatory unitary conductances at this synapse are modelled as alpha functions with an exponential time course of 0.2ms. Synaptic depression with a single exponential recovery is also implemented at this stage, with similar parameters values (*u* = 0.55; *τ* = 25 ms) to the Hodgkin-Huxley-type SBCs.

Each MSO neuron receives 4 (independently-simulated) excitatory cochlear-nucleus inputs from “either ear” in the linear model. Alpha functions are again used to model the unitary excitatory conductances at this synapse, with time-constants: 0.2 ms paired with slow model membranes, and 0.5 ms paired with slow model membranes. Where comparisons of fast and slow model neurons are made, the same adapting auditory nerve inputs are presented to their respective cochlear-nucleus stages.

#### Spike thresholds in the linear-membrane model

The linear-membrane model applies idealised spike thresholds (Remme et al., 2014). A slope threshold (d*v*/dt) is used for all neuronal types. Threshold values for cochlear-nucleus cells are selected to produce good average firing rate without undermining the effects of synaptic depression. Threshold values for the MSO neurons are selected to obtain maximum dynamic range of ITD tuning functions for 200/600 Hz pure tone stimuli (where dynamic range is considered the difference between the maximum and minimum ITD-modulated spike rate). A refractory period of 1 ms is implemented as in Remme et al. (2014) for model SBCs and fast model MSO neurons, whereas a longer refractory period of 2 ms is used for slow model MSO neurons, reflecting longer refractory periods in neurons with lower densities of K_LT_ channels (Rothman & Manis, 2003b, 2003a, 2003c).

#### Calculating lateralisation using d’ values

Lateralisation of a reverberant natural stimulus is judged in the linear model by calculating the mean spike rate difference between a population of 50 ‘left’ and 50 ‘right’ MSO neurons to a single stimulus presentation. d’ values are calculated to quantify the degree of separation of ‘left’ and ‘right’ spike rates using the following equation:

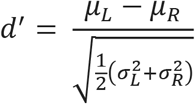

Where *μ*_*L*_ is the mean spike rate for the ‘left’ population of MSO neurons in 5-ms bins; *μ*_*R*_ is the mean spike rate for the ‘right’ population of MSO neurons in 5-ms bins; *σ*_*L*_ is the standard deviation of the ‘left’ MSO population’s spike rate and *σ*_*R*_ is the standard deviation of the ‘right’ population’s spike rate (both calculated in 5-ms bins again).

The sign and magnitude of the d’ allows judgement of whether the linear model lateralises the reverberant stimulus to the correct direction (negative d’ values denote negative ITDs/left of midline; positive d’ values denote positive ITDs/right of midline) and whether this lateralisation judgment is considered significant (d’ > 1 or d’ < −1 indicated a significant lateralisation in the right and left directions respectively).

#### ANOVA and T-test for significant differences in lateralisation

Two-way and three-way ANOVAs are implemented to compare the duration of correct lateralisations (time correctly lateralised = bin duration x number of bins where d’ > 1) by the hemispheric model of SAM tones with non-zero ITDs in either the rising, peak or falling phases as well as the percentage of the phase including a non-zero ITD. A two-tailed T-test is also performed for 200-Hz carrier SAM tones to compare the effect of MSO neuronal type. For the speech stimulus, a paired two-tailed T-test is performed to compare the frequency of incorrect lateralisations (time incorrectly lateralised = bin duration x number of bins where d’ <−1) in the linear model with either adapting or non-adapting auditory nerve inputs.

## Results

### A model MSO neuron driven by adapting SBCs reproduces the frequency-dependent emphasis of ITD information during rising sound-energy

We first explored the extent to which adaptation at model SBCs in the cochlear nuclei can account for the emphasis of ITDs during the rising energy of modulated sounds in a model MSO neuron (Fig. 2B). Acoustic stimuli to the model were amplitude-modulated binaural beats (AMBBs) (Dietz et al., 2013, 2014), in which binaurally presented tones of a slightly different frequency are modulated in amplitude at a rate equal to the difference in frequency across the two ears, and each cycle of rising and falling sound-energy contains a full cycle of interaural phase disparities. AMBBs (Fig. 2A) centred at 600 Hz, with modulation rates 4 to 64 Hz, were presented to a standard model of peripheral auditory processing (Glasberg & Moore, 1990; Zilany et al., 2014), which drove a nonlinear model of brainstem neurons. The model of peripheral auditory processing includes 24 adapting ANFs: twelve each in the left and right ears, with CF 600 Hz. Each of three distinct model ANFs projects an ipsilateral, excitatory synaptic input to one of the eight (four left, and four right) model SBCs. These model synapses uniformly depress (*u* = 0.5) (Rudnicki & Hemmert, 2017) and recover with a 25-ms time-constant (Wang & Manis, 2008). Each model SBC (Rothman & Manis, 2003c) projects a non-depressing, excitatory synaptic input to a multi-compartment model MSO neuron (Brughera et al., 2013) (with minor adjustments described in Methods), for a total of eight excitatory inputs (Couchman et al., 2010). The model SBCs and MSO neuron are Hodgkin-Huxley-type models (Hodgkin & Huxley, 1952) adjusted to temperature 37°C.

Beginning without STP, a nonlinear model including SBCs with very strong, non-depressing, supra-threshold synapses, originally developed for high entrainment (Joris et al., 1994) to account for strong ITD sensitivity in the MSO to unmodulated stimuli (Yin & Chan, 1990), produced only slight adaptation in the spike rates of model SBCs (Fig. 3A, top row). In this condition, model SBCs had excitatory synaptic strength, 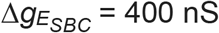; threshold was 34 nS for a single input from rest, using the same model membrane and faster synapses at 38°C (Rothman & Manis, 2003c). With consistently very strong, supra-threshold synapses in SBCs, a model MSO neuron employing relatively weak synapses 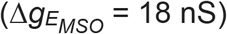 responded strongly to zero ITD, (its pre-determined best-ITD) across the AM cycle: most strongly at peak sound-energy, slightly less strongly during rising energy, and less strongly still during falling energy (Fig. 3A, lower three rows; in a 3-way comparison of spike counts for best-ITD during rising vs. peak vs. falling energy, *P* < 10^−9^ at each modulation frequency). The synchrony index—a measure of temporal alignment of spikes over the AM cycle (see Methods)—ranged from 0.133 at 4 and 64 Hz to 0.161 at 16 Hz for model SBCs, and from 0.595 (64 Hz, best-ITD at peak) to 0.794 (16 Hz, best-ITD during fall) for the model MSO neuron (Rayleigh statistic, *P* < 0.001 in all cases). (AMBB stimuli had increasing IPD (positive beat direction): “best-ITD during rise” denotes start-IPD = 270°; “best-ITD at peak” denotes start-IPD = 180°; and “best-ITD during fall” denotes start-IPD = 90°.)

We next introduced synaptic depression in slightly supra-threshold synapses (maximum 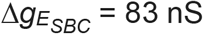) at the model SBCs, which then adapted in their spike rates (Fig. 3B, top row). The model MSO neuron maintains a fixed synaptic strength, 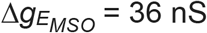, consistent with excitatory fibre conductances of 37 ± 4 nS measured *in vitro* (Couchman et al., 2010). In model SBCs, adaptation in spike rate followed a similar time-course across AM rates, such that at low-to-moderate AM rates (4 to 16 Hz) most of the adaptation occurred by the peak in the AM cycle, and at moderate-to-high rates (16 to 64 Hz) gradual adaptation was evident across much of the AM cycle. Consistent with physiological and behavioural data at 600 Hz (Dietz et al., 2014; Hu et al., 2017), the model MSO neuron responded to its best ITD strongly during rising amplitude, weakly to moderately (and broadly) at peak energy, and only weakly during falling energy (Fig. 3B, lower three rows; in this 3-way comparison of spike counts, *P* ≤ 0.003 at each modulation frequency). Adapting similarly in spike rate to the model SBCs, the model MSO neuron had higher spike counts for best-ITD during rising energy vs. peak energy at 4- to 16-Hz modulation (*P* < 0.02), and during peak vs. falling energy at 16- to 64-Hz modulation (*P* < 0.0025). Synchrony-indices to AM for model SBCs ranged from 0.0485 at 4 Hz to 0.174 at 32 Hz; and for the model MSO neuron, from 0.450 (64 Hz, best-ITD during fall) to 0.814 (64 Hz, best-ITD during rise) (Rayleigh statistic, *P* < 0.01 for model SBCs at 4-Hz AM, otherwise *P* < 0.001).

Consistent with behavioural data at the same 600-Hz sound-frequency, a model including synaptic plasticity at SBCs that in turn drive bilateral inputs to an MSO neuron, responds preferentially to ITDs during the rising energy of sounds. Without this adaptation, the preference for spatial cues during rising sound-energy is absent, suggesting a critical role for monaural input pathways to binaural neurons in emphasising spatial information that is likely to be reliable in reverberant listening conditions.

Mechanisms for sensitivity to ITDs conveyed in the TFS of sounds are generally thought to be consistent across sound-frequencies up to 1400 Hz (higher in some non-human species), covering the range over which ITDs in the TFS are discriminable. Therefore, given the emphasis to ITDs during the rising energy of modulated sounds at 600 Hz—concordant with perceptual and brain-imaging data obtained at a similar frequency (Dietz et al., 2013)—we expected a similar emphasis at the lower sound-frequency of 200 Hz. Nevertheless, despite our model implementing identical elements, we observed a very different pattern of results at 200 Hz compared to 600 Hz. These differences are consistent with behavioural data (Hu et al., 2017): at 600 Hz, human ITD-sensitivity is strongest during rising energy with progressively less ITD sensitivity for peak and falling energy; but at 200 Hz, ITD sensitivity is strong for both peak and rising energy, and again weak for falling energy.

With CFs of the model ANFs now equal to 200 Hz, but otherwise employing the same model with monaural adaptive spike-failures in SBCs, identical to that which emphasised ITDs during the rising energy at 600 Hz, we assessed the relative emphasis of ITD cues in the modulation cycle of a 200-Hz AMBB stimulus, for the modulation rates of 8 and 20 Hz employed by Hu et al. (2017). At 200 Hz, the model SBCs showed slight adaptation in spike rate across the AM cycle (Fig. 4A, top row), less adaptation than at 600 Hz. Spike counts in the model MSO neuron (Fig. 4A, lower three rows) matched the patterns of significance in human ITD-detection for AM stimuli at 200 Hz (Hu et al., 2017): spike counts were slightly higher but not significantly for best-ITD during peak vs. rising energy, and spike counts were significantly different (higher) only for best-ITD at peak energy vs. falling energy (*P* = 0.0079 for modulation at 8 Hz; *P* = 0.0014 for modulation at 20 Hz). Synchrony-indices to AM for model SBCs ranged from 0.0762 at 8 Hz to 0.109 at 20 Hz; and for the model MSO neuron, from 0.433 (20 Hz, best-ITD during fall) to 0.807 (8 Hz, best-ITD during rise) (Rayleigh statistic, *P* < 0.001).

Our nonlinear brainstem model with adapting SBCs reproduces the emphasis of rising energy in ITD encoding by MSO neurons at 600 Hz, and is consistent with the shift from very strong human ITD-sensitivity during rising energy at 600 Hz, to strong sensitivity during both peak and rising energy at 200 Hz, and with relatively weak sensitivity during falling energy at both frequencies.

### A hemispheric-difference model including adapting SBCs correctly lateralises amplitude-modulated stimuli with temporally specific ITDs

To test whether monaural adaptation in combination with a heterogeneous population of MSO neurons (Bondy & Golding, 2018; Remme et al., 2014) is also consistent with behavioural observations (Hu et al., 2017), we employed a neural spiking model of both brain hemispheres, incorporating linear, single-compartment model neurons for computational efficiency in representing the bilateral cochlear nuclei and MSOs (Remme et al., 2014), and generating a neuro-metric measure of lateralisation. Following the model concept of Dietz et al. (2009), two hemispheric channels (‘left’ and ‘right’) comprised distinct MSO model neuron populations (50 neurons each) whose inputs from ‘contralateral’ cochlear nucleus (CN) were delayed by 0.125 cycles of interaural phase difference (IPD, equal to ITD x frequency) (McAlpine et al., 2001) such that they spiked preferentially for sounds arriving from contralateral spatial locations (Fig. 5). The difference in spike rate between hemispheres, calculated (in 5-ms epochs) as a signed d’, was then used as a neuro-metric measure of lateralisation (see Methods) (Devore et al., 2009; McAlpine et al., 2001). A uniform synaptic depression (*u* = 0.55) (Rudnicki & Hemmert, 2017) was implemented independently at each synapse between an ANF and its target SBC of the VCN. Sinusoidally amplitude-modulated (SAM) tones, with non-zero, positive ITDs positioned within the rising, peak, or falling phases of the AM cycle as per Hu et al. (2017), and see above, were presented to the hemispheric-difference model (Fig. 6). As in the behavioural study, the tone frequency (200 Hz or 600 Hz) and the AM rate (8 Hz or 20 Hz) were varied, as was the proportion (20% or 40%) of the AM cycle containing a non-zero ITD.

**Fig. 6.**
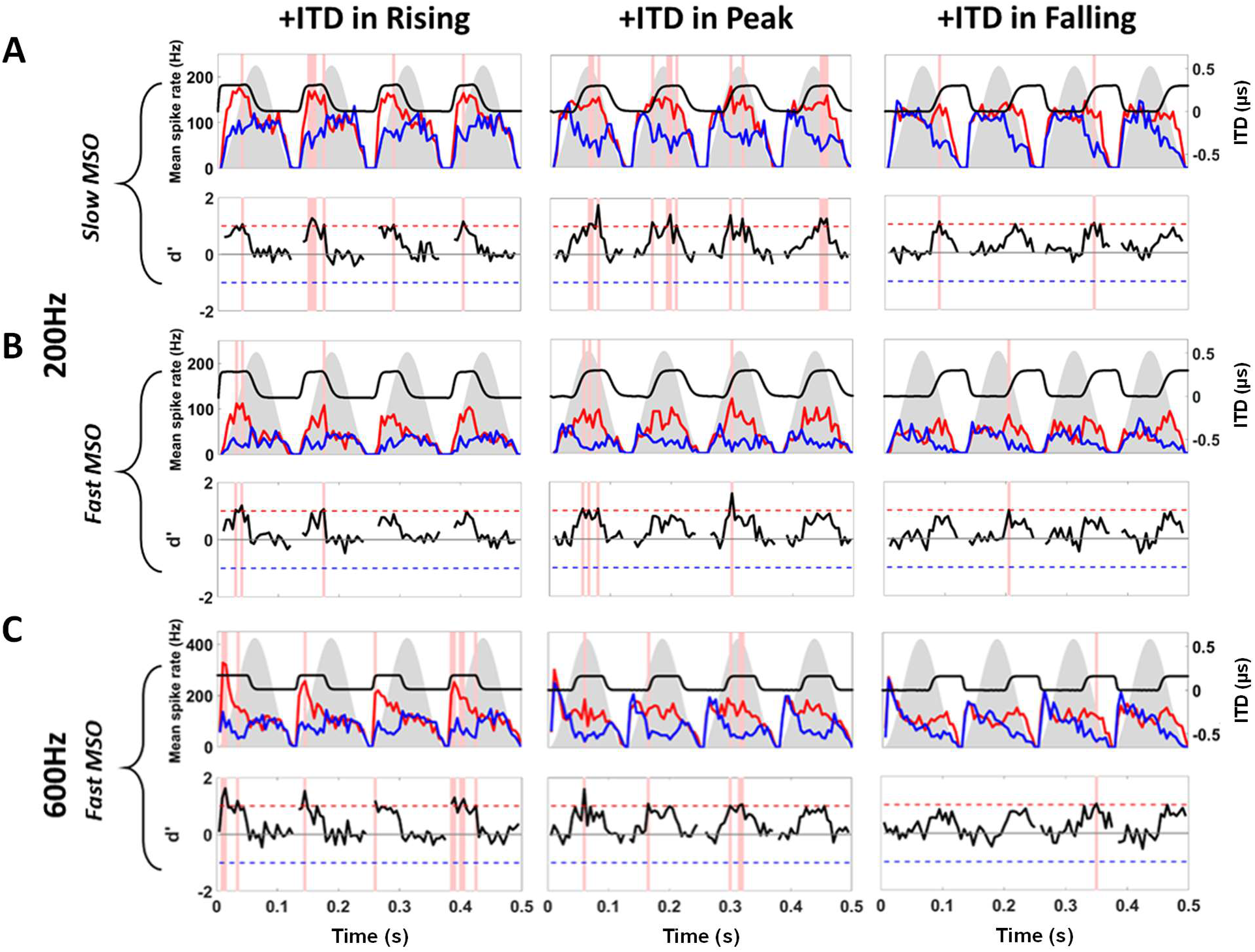
Lateralisation by the hemispheric-difference model at 200 Hz and 600 Hz, each with 8-Hz AM. (A&B) At 200 Hz, both **A** Slow MSO models, and **B** Fast MSO models correctly lateralised +300 μs ITDs (top row, black) to the right, based on d’>1 in any 5ms bin (red vertical bands), most often at peak (middle column) of the AM cycle (grey silhouettes) compared to rising (left column) and falling (right column) phases (Slow MSO: Rising, 7 bins; Peak, 12 bins; Falling, 2 bins. Fast MSO: Rising, 3 bins; Peak, 4 bins; Falling, 1 bin); d’ (bottom rows, black) is a difference in mean spike rates between left (top rows, red) and right (top rows, blue) MSO populations, normalised by variance. The same adapting inputs are used for both MSO neuronal speeds, therefore more correct lateralisations overall by the Slow MSO is a result of its slower integration. **C** At 600 Hz, introducing +150 μs ITD (top row, black) produced strong onset ITD sensitivity in the Fast MSO during rising phase (left column) that decreased across the AM cycle (Fast MSO: Rising, 12 bins; Peak, 5 bins; Falling, 1 bin)

At 600 Hz, model MSO neurons with fast membranes, i.e. akin to the Hodgkin-Huxley-type model, demonstrated a strong sensitivity to the onset ITD, as indicated by the increased number of instances when the AM stimulus containing a (right-leading) +150 μs ITD in its rising-energy phase was correctly lateralised [d’ > 1 in any bin was considered a correct lateralisation of a signal from the ‘right’ (Fig. 6 *pink vertical bands*)]. This ITD sensitivity decreased across the AM cycle reaching a minimum when non-zero ITDs were restricted to the falling-energy phase (Fig. 6 *bottom row*). The trend for onset dominance at 600 Hz was true for both modulation rates (8 Hz and 20 Hz) and when the proportion of the AM cycle containing non-zero ITDs was either 20% or 40% (Fig. 7 *right column*). Notably, in all three portions of the AM cycle, the number of correct lateralisations decreased as the proportion of an AM cycle containing a non-zero ITD was decreased from 40% to 20% (Fig. 7 *right column*, blue).

**Fig. 7.**
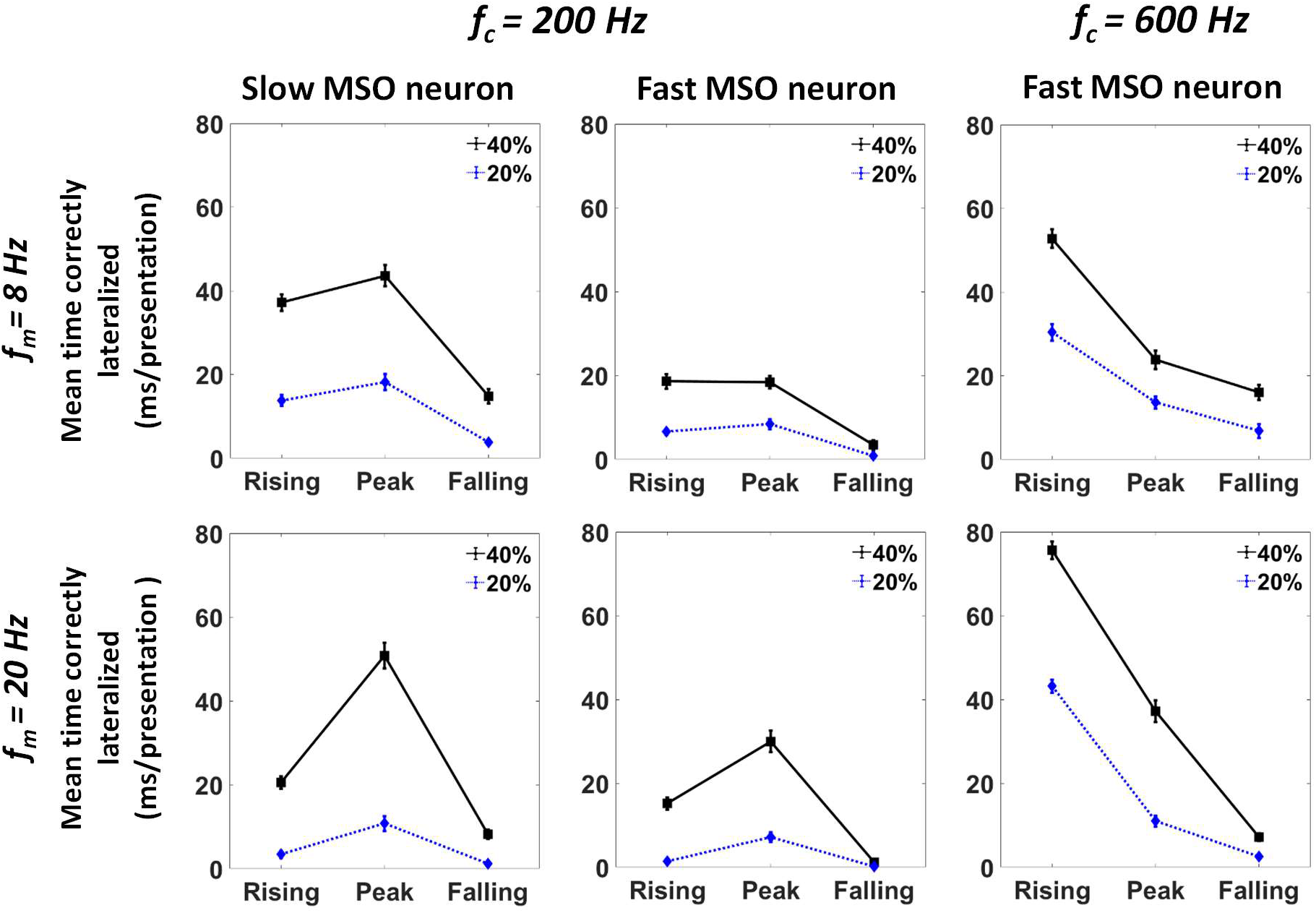
Summarising correct lateralisations of SAM stimuli by the hemispheric-difference model. Mean (±SEM) time correctly lateralised per presentation (number of correct lateralisations * bin duration / 25 presentations) by either Slow MSO neurons at 200 Hz (left column) or Fast MSO neurons at 200 Hz and 600 Hz (middle and right columns respectively), for modulation rates of 8 Hz (top row) and 20 Hz (bottom row). ITDs were inserted into either 20% (blue dotted) or 40% (black solid) of the AM cycle during Rising, Peak, or Falling energy. At 600 Hz, the Fast MSO displays dominant ITD weighting at onset that decreased significantly across AM cycle (2-way ANOVA, interaction of AM cycle phase x Percentage ITD insertion: 8-Hz AM, *F*(2,48) = 8.310, *P* = 0.001; 20-Hz AM, *F*(2,48) = 50.098, *P* = 0.0001). At 200 Hz, the Fast MSO showed equal or better ITD sensitivity at Peak compared to Rising energy (2-way ANOVA, interaction of AM cycle phase x Percentage ITD insertion: 8-Hz AM, *F*(2,48) = 7.83, *P* = 0.001; 20-Hz AM, *F*(2,48) = 36.01, *P* = 0.0001). Slow MSO generated more correct lateralisations than Fast MSO (2-way ANOVA, interaction of AM cycle phase x Neuron Type: 8-Hz AM, *F*(2,48) = 18.82, *P* = 0.0001; 20-Hz AM, *F*(2,48) = 13.12, *P* = 0.0001) with equal or augmented ITD sensitivity at Peak for 40% non-zero ITD (Paired two-tailed T-test, Mean Peak – Rising, N = 25: 8-Hz AM, *t*(24) = 2.21, *P* = 0.037; 20-Hz AM, *t*(24) = 2.21, *P* = 2.37 × 10^−4^). Correct lateralisations also increased as the percentage of AM phase containing non-zero ITD was raised from 20% to 40% (3-way ANOVA, main effect of Percentage ITD insertion: Fast MSO, *F*(1,24) = 418.78, *P* = 0.0001; Slow MSO, *F*(1,24) = 285.622, *P* = 0.0001)

When an amplitude-modulated 200-Hz tone (with a right-leading, non-zero ITD of +300 μs) was presented to the fast MSO model, the onset dominance observed at 600 Hz was replaced by an increased weighting of ITD cues towards the peak of the AM cycle (Fig. 6). Indeed, the frequency of correct lateralisations at the peak of AM cycles was either equal to (8-Hz AM stimuli) or higher (20-Hz AM stimuli) than that observed at corresponding onset phases (Fig. 7 *middle column*). As at 600 Hz, reducing the proportion of the AM cycle containing a non-zero ITD from 40% to 20% also generated fewer correct lateralisations across all AM phases (Fig. 7 *middle column*, blue). Although the fast MSO model could reproduce behavioural trends (Hu et al., 2017) at both carrier frequencies, it generated fewer correct lateralisations overall at 200 Hz (Fig. 7 *middle* and *right columns*). We therefore presented the same 200-Hz carrier stimulus to the linear model MSO neurons with slower membrane properties (Fig. 6). This generated a higher number of correct lateralisations overall, while maintaining, or even augmenting, the maximum weighting of ITD at the peak AM energy in all conditions tested (Fig. 7 *left column*).

Our hemispheric-difference model, therefore, suggests that a slower, more integrative MSO neuron may assist lateralisation in lower-frequency channels, in particular by extending the extraction of ITD information towards peak energy in the AM cycle, where human ITD detection is best when comparing at 200 Hz across the AM cycle (Hu et al., 2017). Additionally, our model qualitatively matched the behavioural data by demonstrating a higher propensity to lateralise stimuli of either frequency when 40% vs. 20% of the AM cycle contained a non-zero ITD.

### Adapting SBCs improve the ability of a hemispheric-difference model to lateralise speech in artificial reverberation

If neural adaptation prior to binaural integration contributes to accurate source localisation, we would expect it to aid lateralisation of speech in a reverberant room, by emphasising ITD cues associated with direct sound over reverberation (Fig. 1 *far left*) (Dietz et al., 2014). To test this, speech signals were presented to the neural model of lateralisation under anechoic (non-reverberant) conditions, and simulated reverberant conditions with direct sound and early reflections off virtual walls (Dietz et al., 2014). We then assessed whether the model correctly lateralised the side to which the source was located in the presence or absence of synaptic depression in SBCs of the cochlear nuclei.

In the anechoic condition (Fig. 8A), a word token (‘church’) was presented to the non-adapting hemispheric-difference model from a virtual location 1 metre from the target and 30 degrees to the right of the midline. The speech waveform—gammatone-filtered at 600 Hz (Fig. 8A *top row*)—was more intense at the right ear (red), and led the left-ear signal (blue) by an average ITD of +363 μs (Fig. 8A *top row*, dark grey). This generated a larger response from model neurons in the ‘left’ MSO (Fig. 8A *middle row*) and multiple bins throughout the speech token where d’ values were > 1 (Fig. 8A *pink vertical bands*), indicating that the talker was correctly lateralised to the right in the non-adapting hemispheric-difference model.

In the reverberant condition, the same word token was presented to the non-adapting hemispheric-difference model from the same angle and distance, however it was also followed by two delayed copies generated by early reflections off virtual walls added behind, and to the left of, the listener (Fig. 1B *far left*). This mixture of sound arriving direct from the source and early reflective copies led to a more complex waveform (including a more intense signal at the ‘left ear’ (direct-to-reverberant ratio: −4dB; mean ILD: −3 dB)) whose running ITDs fluctuated between extremely large positive and negative values, culminating in an average IPD of +155°, a value that can easily lead to wrong-side lateralization for sine-tones (Yost, 1981). Although the resulting d’ output exceeded 1 in eleven epochs coinciding with waveform onsets (Fig. 8B, *left*, *pink vertical bands*), it was also briefly lower than −1 in the opposite direction during three intermediate epochs (Fig. 8B, *left*, *blue vertical bands*), indicating a potentially ambiguous left-right localisation output.

We then introduced synaptic depression to the monaurally driven inputs from ANFs to the cochlear nuclei of the hemispheric-difference model, and presented the same reverberant stimulus (Fig. 8B). Despite the potentially confounding ITD (and ILD) cues (Fig. 8B *top row*), adding adaptation in the monaural input pathways enhanced the performance of the binaural model MSO neurons, and generated correct lateralisations of the true source to the right: the d’ output exceeding 1 in six time bins without ever crossing below −1 (Fig. 8B *bottom row, right*). When the number of incorrect lateralisations was tallied over a hundred presentations of the reverberant stimulus (Fig. 8C), incorrect lateralisations were observed five times less frequently when synaptically depressing auditory nerve inputs were included in the hemispheric-difference model (mean incorrectly lateralised time for non-depressing synaptic inputs = 6.75 ± 0.50 ms/presentation vs. mean incorrectly lateralised time for depressing synaptic inputs = 1.3 ± 0.24 ms/presentation). This neuro-metric profile agrees with the psychoacoustic percept as has been previously described for similar word tokens (Dietz et al., 2014), suggesting that monaural adaptation increases the weighting of spatial information during rising sound-energy to improve lateralisation of ethological sounds in reverberant environments.

## Discussion

### The adapting brainstem suppresses responses to late-arriving reverberant sound while encoding spatial cues

Localising sound-sources in reverberant environments is critical to prey and predator alike, and improves communication in human listeners. Despite reflections from acoustically opaque surfaces degrading spatial cues, localisation must remain accurate. Hypothesising that adaptive brainstem mechanisms suppress responses to late-arriving reverberant sound, thus emphasising early-arriving sound direct from the source for the encoding of spatial cues, we computationally explored the contribution of monaural brainstem adaption to binaural sound-source lateralisation. In our models, accurate lateralisation in reverberation is enhanced by adapting SBCs in the VCN, which project bilaterally to the MSO, a site of primary binaural integration. We show that pre-binaural adaptation can account for the observed ability of MSO neurons to respond preferentially to ITDs conveyed during the early, rising-energy portion of low-frequency sounds near 600 Hz (Dietz et al., 2014). Reflecting perception of normal-hearing listeners at 500 Hz (Dietz et al., 2013), our models successfully glimpse TFS-ITD cues conveyed during the AM cycle’s rising portion, suppressing spatial information conveyed in all later portions, including that with highest energy.

### Pre-binaural adaptation with appropriate temporal properties supported the reproduction of in vivo MSO responses

Our model MSO neurons reproduced *in vivo* responses to AMBBs (Dietz et al., 2014), using pre-binaural adaptation at model ANFs (Zilany et al., 2014, 2009) and SBCs. Adaptive effects of glycinergic inhibition at SBCs *in vivo* (Keine & Rübsamen, 2015; Keine et al., 2016; Kuenzel et al., 2011, 2015) are phenomena-logically modelled using STP as measured *in vitro* at calyceal synapses from ANFs to SBCs (Oleskevich et al., 2000; Wang & Manis, 2008; Yang & Xu-Friedman, 2009, 2015), noting their nearly identical temporal properties: the time-constant of decay in glycinergic inhibition, 23.9 ms (Kuenzel et al., 2015); and the time-constant of recovery in STP, 25 ms (Wang & Manis, 2008). Acknowledging the disputed role of STP, its similar time-course for adaptation compared with glycinergic inhibition supported modelling of *in vivo* MSO responses to AMBB stimuli.

Actual STP *in vivo* is reportedly weak due to GABAB-receptor-mediated limitation of pre-synaptic vesicle release, maintaining initially weaker but more consistent synapses, including slightly supra-threshold calyces (Chanda & Xu-Friedman, 2010; Keine et al., 2016; Kuenzel et al., 2011; Lorteije et al., 2009). Accordingly, STP was not required at model MSO neurons. Significant adaptation on a 25-ms time-scale is not expected to originate at MSO neurons *in vivo*: glycinergic inhibition is faster (time-constant, 2 ms) (Magnusson et al., 2005); GABA presumably limits STP, and GABA_B_-receptor-mediated synaptic adaptation is slower (time-scale, 500 ms) (Stange et al., 2013).

Inhibition at SBCs may not entirely explain pre-binaural adaptation: *in vivo,* some bushy cells (BCs) adapt similarly with and without pharmacological blocking of either glycinergic or GABA_A_ inhibition (Gai and Carney, 2008). Spike-rate adaptation in ANFs (Zilany and Carney, 2010) can also temporally shape monaural inputs, as suggested by examples of increasing onset-emphasis from adapting model ANFs to globular BCs, for AM rates up to 64Hz (Stecker, 2020). Number, synaptic strength, and CF-span of ANF inputs to BCs likely influence how ANF adaptation contributes to pre-binaural adaptation overall (Ashida et al., 2019; Brughera et al., 1996; Carney, 1990, 1992; Rudnicki & Hemmert, 2017).

### Frequency-dependent emphasis of early-arriving sound reflects natural frequency-profiles in reverberant energy

Natural outdoor acoustics have seemingly influenced brain mechanisms that suppress responses to reverberation. In many outdoor environments, including forests, fields, and streets, reverberation-time (‘T60’—the time for reverberant energy to fall by 60 decibels) decreases as sound-frequency decreases below 1500 Hz (Traer & McDermott, 2016). Reflecting this frequency-profile in reverberation-time, our models are consistent with the frequency-dependent behavioural emphasis of ITD during rising and peak sound-energy (Hu et al., 2017), by increasing the weighting of peak energy with decreasing low frequency where reverberation is less energetic. Applying identical parameter values in our Hodgkin-Huxley-type model MSO neuron as sound-frequency decreased from 600 to 200 Hz, model neurons transitioned from responding preferentially to ITDs during rising energy at 600 Hz, to responding equally strongly to ITDs during rising and peak energy at 200 Hz. At both frequencies, model neurons aptly responded only weakly to ITD during falling energy, when direct sound is most likely to be corrupted by reverberation. This is consistent with listening behaviour: at 600 Hz, human listeners are more sensitive to ITDs conveyed during rising-energy than to ITDs at the energy peak; at 200 Hz, listeners are equally sensitive to ITDs conveyed during rising and peak energy; listeners are least sensitive to ITD during falling sound-energy at both sound-frequencies. These data, and our models, suggest that spatial auditory brain mechanisms transmit reliable information, and suppress unreliable information, accounting for natural frequency-profiles in reverberant energy.

At very low frequencies, including 200 Hz, where reverberant energy in natural, outdoor scenes is low, suppression of spatial information during peak energy is less important. However, in modern, indoor listening environments, characterised by enclosures with highly reflective walls, reverberation can be high even at very low frequencies. Indoor reverberation, weakly suppressing neurons, and increasing perceptual thresholds for ITD at sound-frequencies below 500 Hz (Brughera et al., 2013), may all contribute to the observed low perceptual weighting of very low frequencies when localising broadband sound (Ihlefeld & Shinn-Cunningham, 2011). This consistent perceptual down-weighting for localisation, presumably by brain centres above the brainstem, occurs even as these very low sound-frequencies contain and apparently convey vital speech information, including the fundamental frequency, and first formant of vowels.

Contrasting the weak suppression of late-arriving sound and weak reverberation at 200 Hz, with the strong suppression at 600 Hz and increasing reverberation with higher sound-frequency up to 1500 Hz, suggests the possibility that the human brainstem effectively suppresses responses to reverberation for sound-frequencies from 500 to 1200 Hz, a frequency range that despite being relatively high in reverberation produces the lowest perceptual thresholds for ITD discrimination in human listeners (Brughera et al., 2013; Klumpp & Eady, 1956), and dominates the percept of auditory spatial cues (Ihlefeld & Shinn-Cunningham, 2011; Shinn-Cunningham et al., 1995; Xia et al., 2010).

### Model MSO neurons with a plausible range of membrane speeds are effective at low sound-frequencies

MSO neurons with slower intrinsic properties than typically recorded in principal MSO neurons (though faster than other types of neurons in the central nervous system) indicate some degree of heterogeneity in the nucleus (Bondy & Golding, 2018; Remme et al., 2014). By not expressing more ion channels than required, the slower MSO neurons encode ITD efficiently, realising a cellular energetic advantage. Introducing moderately slow MSO membranes (labelled “Slow MSO”) boosted the hemispheric-difference model’s ability to lateralise 200-Hz signals, especially at their energetic peak, suggesting a functional benefit of these slower membranes at lower frequencies.

### Adapting SBCs enhance correct lateralisation of reverberant speech

Early reflections—those following the direct signal by 50 ms or less (Bradley et al., 2003)—can disrupt low-frequency sound localisation by conveying spurious TFS-ITD cues (Gourévitch & Brette, 2012). Yet normal-hearing listeners can locate sound-sources, including talkers in reverberant environments (Bregman, 1997). By demonstrating that within a hemispheric 2-channel model, the addition of SBC adaptation emphasises ITD cues in sound onsets, our data suggest one means by which complex sounds, including speech, can be reliably lateralised.

Whilst our adapting hemispheric-difference model correctly lateralised reverberant speech, this was based on its d’ output surpassing a threshold value in only six of the ninety 5-ms bins with a correct ITD cue available. Although this neuro-metric measure may not appear particularly robust, it should be remembered that only the 600-Hz frequency channel was examined for speech lateralisation. Localisation in reverberant conditions and the precedence effect—the suppression of spatial information for late-arriving sounds—both demonstrate strong weighting of cues from 500 to 750 Hz (Ihlefeld & Shinn-Cunningham, 2011; Shinn-Cunningham et al., 1995; Xia et al., 2010), but they also highlight subjects’ inability to localise pure tones in echoic conditions, suggesting that enhanced speech localisation in reverberant conditions involves localisation cues across the low-frequency spectrum. Adding a midbrain stage, where inferior colliculus (IC) neurons receive inputs from MSO neurons across a wide range of CFs, would further test the extent to which adaptation in the VCN improves speech localisation in reverberant environments.

### Post-binaural adaptation

Onset-cue dominance is demonstrated extensively in the precedence effect (Brown et al., 2015; Wallach et al., 1949), where suppression of neural responses to a brief lagging stimulus occurs over a range of delays between leading and lagging stimuli. Future localisation models might involve monaural parallel inhibition that is feedforward from ANFs to dorsal cochlear nucleus (DCN), with the relatively complex DCN inhibiting the VCN (Keine et al., 2017; Zheng & Voigt, 2006a, 2006b). A broader model with these adaptive brainstem elements, combined with additional inhibitory echo-suppressive mechanisms, such as delayed inhibition to the midbrain (Burger & Pollak, 2001; Kidd & Kelly, 1996; Pecka et al., 2007), might explore whether these mechanisms act cooperatively in robust onset-ITD processing in reverberant conditions. Brainstem and midbrain mechanisms may combine additively or act independently for different stimuli. At least for ongoing, amplitude-modulated sounds, the emphasis of early-arriving sound in ITD-encoding by MSO neurons, combined with the observed lack of increased emphasis at the IC (Dietz et al., 2014), suggests that brainstem nuclei contribute significantly to sound-source localisation in reverberation.

### Early-arriving spatial cues for bilateral cochlear-implant (bCI) listeners

Although listeners with bCIs are most sensitive to ITD during peaks in sound-energy (Hu et al., 2017), acoustically the most accurate ITD-information for source location occurs during rising sound-energy (Dietz et al., 2013). To maximise ITD sensitivity, and provide spatial information that emphasises sound-sources, bCI processors can, during peak energy, provide pulse bursts that overcome adaptation (Srinivasan et al., 2020, 2018) to convey spatial information derived milliseconds earlier during rising energy, from a calculation triggered by the preceding energy-minimum and subsequent energy-increase. Rapidly updating binaural masks (Cantu, 2018) that enhance target sound-sources while preserving spatial cues can also be applied.

## Conclusions

Our models suggest that adaptive brainstem mechanisms contribute to sound-source localisation, emphasising early-arriving sound which is relatively high in direct sound that conveys reliable spatial information during neural encoding, by suppressing responses to late-arriving sound which is relatively high in reverberation. The frequency-dependent emphasis of auditory spatial information conveyed in early-arriving sound is consistent with brain mechanisms that transmit reliable information, and suppress unreliable information. As the auditory brainstem encodes ITDs for determining sound-source locations, its suppression of late-arriving spatial information promotes accuracy and accounts for typical frequency-profiles of reverberant energy in natural outdoor scenes.

## Declarations

### Funding

This work was supported by Australian Research Council Laureate Fellowship (FL160100108) awarded to David McAlpine

### Conflicts of interest/Competing interests

The authors declare that they have no conflict of interest, and no competing interest.

### Ethics approval

For this strictly computational study, no approval was required.

### Consent to participate

For this strictly computational study, no consent was required.

### Consent for publication

For this strictly computational study, there are no human subjects from whom consent is required. All authors and responsible authorities approve the publication of this manuscript.

### Availability of data and material

For the nonlinear model: data, analysis scripts, and code are available at figshare: https://doi.org/10.6084/m9.figshare.11955219.v1

For the linear model: data, analysis scripts, and code are available at figshare: https://doi.org/10.6084/m9.figshare.9899018.v1

### Code availability

For the nonlinear model: code is available at Github: https://github.com/AndrewBrughera/Mso_SbcStp_EE1

For the linear model: code is available at figshare: https://doi.org/10.6084/m9.figshare.9899018.v1

### Author Contributions

Andrew Brughera & Jason Mikiel-Hunter contributed equally.

**Conceptualisation:** David McAlpine & Mathias Dietz

**Data Curation:** Jason Mikiel-Hunter & Andrew Brughera

**Formal Analysis:** Jason Mikiel-Hunter & Andrew Brughera

**Funding Acquisition:** David McAlpine

**Investigation:** Andrew Brughera & Jason Mikiel-Hunter

**Methodology:** Andrew Brughera (nonlinear modelling) & Jason Mikiel-Hunter (linear modelling)

**Resources:** David McAlpine, Andrew Brughera, & Jason Mikiel-Hunter

**Software:** Andrew Brughera & Jason Mikiel-Hunter

**Supervision:** David McAlpine

**Visualisation:** Jason Mikiel-Hunter & Andrew Brughera

**Validation:** Andrew Brughera & Jason Mikiel-Hunter

**Writing – Original Draft Preparation:** Andrew Brughera, Jason Mikiel-Hunter, & David McAlpine

**Writing – Review & Editing:** Andrew Brughera, David McAlpine, Mathias Dietz, & Jason Mikiel-Hunter

---

**Supplemental Fig. S1.**
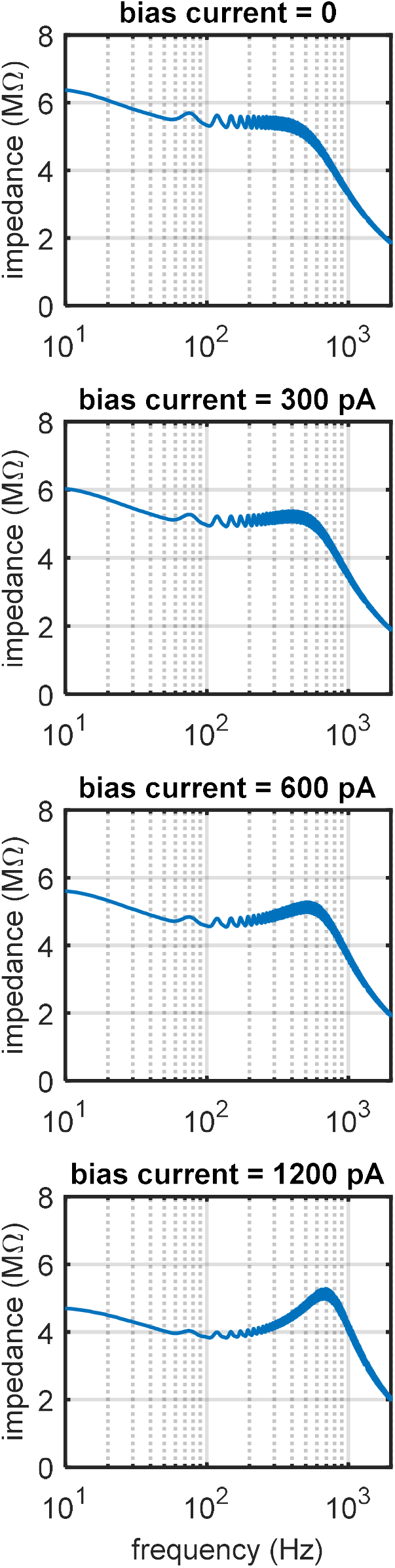
Somatic membrane impedance magnitude as a function of frequency in the Hodgkin-Huxley-type model MSO neuron. Injected membrane current is a steady bias current, plus a frequency sweep in a sine wave with peak amplitude 250 pA. A resonance in membrane impedance emergences with increases in bias current (*I*_*BIAS*_) and resulting increases in membrane holding potential (*V*_*HOLD*_): **A** *I*_*BIAS*_ 0 pA, *V*_*HOLD*_ −60.52 mV, no resonance; **B** *I*_*BIAS*_ 300 pA, *V*_*HOLD*_ −59.22 mV, resonance frequency, f_0_ = 408 Hz; **C** *I*_*BIAS*_ 600 pA, *V*_*HOLD*_ −57.88 mV, f_0_ = 513 Hz; **D** *I*_*BIAS*_ 1200 pA, *V*_*HOLD*_ −55.24 mV, f_0_ = 692 Hz. In **B**, slightly above resting potential, the resonance at 408 Hz indicates a membrane time-constant of 1/(2π × 408 Hz) = 0.390 ms.

## Notes

#### Summary of Updates

Added and revised analysis and discussion of adaptive mechanisms, including short-term plasticity and inhibition with references. Removed unsupported conclusions concerning Kv1 channels. These revisions affect all sections. Methods of the nonlinear model have added connection with physiological references, and a supplemental figure on membrane impedance.

## References

Ashida G, Heinermann HT, & Kretzberg J (2019) Neuronal population model of globular bushy cells covering unit-to-unit variability. PLoS Comput Biol 15(12):1–38. https://doi.org/10.1371/journal.pcbi.1007563

Bondy B, & Golding NL (2018) Variations in intrinsic and synaptic properties in MSO neurons confer a spectrum of ITD sensitivity. ARO Midwinter Meeting, San Diego, CA, USA.

Bradley JS, Sato H, & Picard M (2003) On the importance of early reflections for speech in rooms. J Acoust Soc Am 113(6):3233. https://doi.org/10.1121/1.1570439

Bregman AS (1997) Auditory Scene Analysis - The Perceptual Organization of Sound. Cambridge, Massachusetts, U.S.A.: M.I.T. Press.

Brown AD, Jones HG, Kan A, Thakkar T, Stecker GC, Goupell MJ, & Litovsky RY (2015) Evidence for a neural source of the precedence effect in sound localization. J Neurophysiol 114(5):2991–3001. https://doi.org/10.1152/jn.00243.2015

Brughera A, Dunai L, & Hartmann WM (2013) Human interaural time difference thresholds for sine tones: The high-frequency limit. J Acoust Soc Am 133(5):2839–2855. https://doi.org/10.1121/1.4795778

Brughera AR, Stutman ER, Carney LH, & Colburn HS (1996) A Model with Excitation and Inhibition for Cells in the Medial Superior Olive. Audit Neurosci 2:219–233. Retrieved from https://www.urmc.rochester.edu/MediaLibraries/URMCMedia/labs/carney-lab/codes/Brughera-AuditoryNeuro-1996.pdf

Burger RM, & Pollak GD (2001) Reversible Inactivation of the Dorsal Nucleus of the Lateral Lemniscus Reveals Its Role in the Processing of Multiple Sound Sources in the Inferior Colliculus of Bats. J Neurosci 21(13):4830–4843. https://doi.org/10.1523/jneurosci.21-13-04830.2001

Cantu M (2018) Sound source segregation of multiple concurrent talkers via short-time target cancellation. Ph.D. dissertation, Boston University. Retrieved from https://open.bu.edu/handle/2144/32082

Carney LH (1990) Sensitivities of cells in anteroventral cochlear nucleus of cat to spatiotemporal discharge patterns across primary afferents. J Neurophysiol 64(2):437–456. https://doi.org/10.1152/jn.1990.64.2.437

Carney LH (1992) Modelling the sensitivity of cells in the anteroventral cochlear nucleus to spatiotemporal discharge patterns. Philos Trans R Soc Lond B Biol Sci 336(1278):403–406. https://doi.org/10.1098/rstb.1992.0075

Chanda S, & Xu-Friedman MA (2010) Neuromodulation by GABA Converts a Relay Into a Coincidence Detector. J Neurophysiol 104(4):2063–2074. https://doi.org/10.1152/jn.00474.2010

Cherry EC (1953) Some experiments on the recognition of speech, with one and with 2 ears. JASA (25):975–979. https://doi.org/10.1121/1.1907229

Couchman K, Grothe B, & Felmy F (2010) Medial Superior Olivary Neurons Receive Surprisingly Few Excitatory and Inhibitory Inputs with Balanced Strength and Short-Term Dynamics. J Neurosci 30(50):17111–17121. https://doi.org/10.1523/JNEUROSCI.1760-10.2010

Devore S, Ihlefeld A, Hancock K, Shinn-Cunningham B, & Delgutte B (2009) Accurate Sound Localization in Reverberant Environments Is Mediated by Robust Encoding of Spatial Cues in the Auditory Midbrain. Neuron 62(1):123–134. https://doi.org/10.1016/j.neuron.2009.02.018

Dietz M, Ewert SD, & Hohmann V (2009) Model-based direction estimation of concurrent speakers in the horizontal plane. J Acoust Soc Am 125(4):2527. https://doi.org/10.1121/1.4783519

Dietz M, Marquardt T, Salminen NH, & McAlpine D (2013) Emphasis of spatial cues in the temporal fine structure during the rising segments of amplitude-modulated sounds. Proc Natl Acad Sci 110(37):15151–15156. https://doi.org/10.1073/pnas.1309712110

Dietz M, Marquardt T, Stange A, Pecka M, Grothe B, & McAlpine D (2014) Emphasis of spatial cues in the temporal fine structure during the rising segments of amplitude-modulated sounds II: single-neuron recordings. J Neurophysiol 111(10):1973–1985. https://doi.org/10.1152/jn.00681.2013

Fischl MJ, Combs TD, Klug A, Grothe B, & Burger RM (2012) Modulation of synaptic input by GABAB receptors improves coincidence detection for computation of sound location. J Physiol 590(13):3047–3066. https://doi.org/10.1113/jphysiol.2011.226233

Fitzpatrick DC, Kuwada S, Kim DO, Parham K, & Batra R (1999) Responses of neurons to click-pairs as simulated echoes: Auditory nerve to auditory cortex. J Acoust Soc Am 106(6):3460–3472. https://doi.org/10.1121/1.428199

Franken TP, Roberts MT, Wei L, Golding NL, & Joris PX (2015) In vivo coincidence detection in mammalian sound localization generates phase delays. Nat Neurosci 18(3):444–454. https://doi.org/10.1038/nn.3948

Glasberg BR, & Moore BCJ (1990) Derivation of auditory filter shapes from notched-noise data. Hear Res 47:103–138. https://doi.org/10.1016/0378-5955(90)90170-T

Gourévitch B, & Brette R (2012) The impact of early reflections on binaural cues. J Acoust Soc Am 132(1):9–27. https://doi.org/10.1121/1.4726052

Haas H (1951) The influence of a single echo on the audibility of speech. Acoustica 1(2):49–58. Retrieved from https://www.ingentaconnect.com/content/dav/aaua/1951/00000001/00000002/art00003

Hodgkin AL, & Huxley AF (1952) A quantitative description of ion currents and its applications to conduction and excitation in nerve membranes. J Physiol 117:500–544. https://doi.org/10.1113/jphysiol.1952.sp004764

Hu H, Ewert SD, McAlpine D, & Dietz M (2017) Differences in the temporal course of interaural time difference sensitivity between acoustic and electric hearing in amplitude modulated stimuli. J Acoust Soc Am 141(3):1862–1873. https://doi.org/10.1121/1.4977014

Hutcheon B, & Yarom Y (2000) Resonance, oscillation and the intrinsic. Trends Neurosci 23(5):216–222. https://doi.org/10.1016/S0166-2236(00)01547-2

Ihlefeld A, & Shinn-Cunningham BG (2011) Effect of source spectrum on sound localization in an everyday reverberant room. J Acoust Soc Am 130(1):324–333. https://doi.org/10.1121/1.3596476

Johnson DH (1980) The relationship between spike rate and synchrony in responses of auditory-nerve fibers to single tones. J Acoust Soc Am 68:1115–1122. https://doi.org/10.1121/1.384982

Joris PX, Smith PH, Yin TC, & Carney LH (1994) Enhancement of neural synchronization in the anteroventral cochlear nucleus. II. Responses in the tuning curve tail. J Neurophysiol. https://doi.org/10.1152/jn.1994.71.3.1037

Joris PX, & Yin TCT (1992) Responses to amplitude-modulated tones in the auditory nerve of the cat. J Acoust Soc Am 91:215–232. https://doi.org/10.1121/1.402757

Keine C, & Rübsamen R (2015) Inhibition shapes acoustic responsiveness in spherical bushy cells. J Neurosci 35(22):8579–8592. https://doi.org/10.1523/JNEUROSCI.0133-15.2015

Keine C, Rübsamen R, & Englitz B (2016) Inhibition in the auditory brainstem enhances signal representation and regulates gain in complex acoustic environments. ELife 5:1–33. https://doi.org/10.7554/eLife.19295

Keine C, Rübsamen R, & Englitz B (2017) Signal integration at spherical bushy cells enhances representation of temporal structure but limits its range. ELife 6:1–16. https://doi.org/10.7554/eLife.29639

Kidd SA, & Kelly JB (1996) Contribution of the Dorsal Nucleus of the Lateral Lemniscus to Binaural Responses in the Inferior Colliculus of the Rat: Interaural Time Delays. J Neurosci 16(22):7390–7397. https://doi.org/10.1523/jneurosci.16-22-07390.1996

Klumpp RG, & Eady HR (1956) Some Measurements of Interaural Time Difference Thresholds. J Acoust Soc Am 28(5):859–860. https://doi.org/10.1121/1.1908493

Kuenzel T, Borst JGG, & van der Heijden M (2011) Factors Controlling the Input-Output Relationship of Spherical Bushy Cells in the Gerbil Cochlear Nucleus. J Neurosci 31(11):4260–4273. https://doi.org/10.1523/JNEUROSCI.5433-10.2011

Kuenzel T, Nerlich J, Wagner H, Rubsamen R, & Milenkovic I (2015) Inhibitory properties underlying non-monotonic input-output relationship in low-frequency spherical bushy neurons of the gerbil. Front Neural Circuits 9(March):1–14. https://doi.org/10.3389/fncir.2015.00014

Lehnert S, Ford MC, Alexandrova O, Hellmundt F, Felmy F, Grothe B, & Leibold C (2014) Action Potential Generation in an Anatomically Constrained Model of Medial Superior Olive Axons. J Neurosci 34(15):5370–5384. https://doi.org/10.1523/JNEUROSCI.4038-13.2014

Liebenthal E, & Pratt H (2002) Human auditory cortex electrophysiological correlates of the precedence effect: Binaural echo lateralization suppression. J Acoust Soc Am 106(1):291–303. https://doi.org/10.1121/1.427057

Litovsky RY, & Yin TC (1998a) Physiological studies of the precedence effect in the inferior colliculus of the cat. I. Correlates of psychophysics. J Neurophysiol 80(3):1285–1301. https://doi.org/10.1121/1.423072

Litovsky RY, & Yin TCT (1998b) Physiological Studies of the Precedence Effect in the Inferior Colliculus of the Cat. II. Neural Mechanisms. J Neurophysiol 80(3):1302–1316. https://doi.org/10.1152/jn.1998.80.3.1302

Lorente De No R (1981) The Primary Acoustic Nuclei. New York: Raven.

Lorteije JAM, Rusu SI, Kushmerick C, & Borst JGG (2009) Reliability and precision of the mouse calyx of Held synapse. J Neurosci 29(44):13770–13784. https://doi.org/10.1523/JNEUROSCI.3285-09.2009

Magnusson AK, Kapfer C, Grothe B, & Koch U (2005) Maturation of glycinergic inhibition in the gerbil medial superior olive after hearing onset. J Physiol 568(2):497–512. https://doi.org/10.1113/jphysiol.2005.094763

Mathews PJ, Jercog PE, Rinzel J, Scott LL, & Golding NL (2010) Control of submillisecond synaptic timing in binaural coincidence detectors by K v 1 channels. Nat Neurosci 13(5):601–609. https://doi.org/10.1038/nn.2530

McAlpine D, Jiang D, & Palmer AR (2001) A neural code for low-frequency sound localization in mammals. Nat Neurosci 4(4):396–401. https://doi.org/10.1038/86049

Moser T, & Beutner D (2000) Kinetics of exocytosis and endocytosis at the cochlear inner hair cell afferent synapse of the mouse. Proc Natl Acad Sci 97(1):883–888. https://doi.org/10.1073/pnas.97.2.883

Nilsson JW, & Riedel SA (2008) Electric Circuits. Prentice Hall.

Oleskevich S, Clements J, & Walmsley B (2000) Release probability modulates short-term plasticity at a rat giant terminal. J Physiol 524(2):513–523. https://doi.org/10.1111/j.1469-7793.2000.00513.x

Pecka M, Zahn TP, Saunier-Rebori B, Siveke I, Felmy F, Wiegrebe L, … Grothe B (2007) Inhibiting the Inhibition: A Neuronal Network for Sound Localization in Reverberant Environments. J Neurosci 27(7):1782–1790. https://doi.org/10.1523/JNEUROSCI.5335-06.2007

Puil E, Gimbarzevsky B, & Miura RM (1986) Quantification of membrane properties of trigeminal root ganglion neurons in guinea pigs. J Neurophysiol 55(5):995–1016. https://doi.org/10.1152/jn.1986.55.5.995

Remme MWH, Donato R, Mikiel-Hunter J, Ballestero JA, Foster S, Rinzel J, & McAlpine D (2014) Subthreshold resonance properties contribute to the efficient coding of auditory spatial cues. Proc Natl Acad Sci 111(22):E2339–E2348. https://doi.org/10.1073/pnas.1316216111

Rhode WS (1976) A test for the significance of the mean direction and the concentration parameter of a circular distribution. Madison Univ Wisconsin Dept Neurophysiol Report. Retrieved from http://www.neurophys.wisc.edu/comp/docs/not011/not011.html

Rhode WS, & Greenberg S (1994) Encoding of amplitude modulation in the cochlear nucleus of the cat. J Neurophysiol 71(5):1797–1825. https://doi.org/10.1152/jn.1994.71.5.1797

Rothman JS, & Manis PB (2003a) Differential Expression of Three Distinct Potassium Currents in the Ventral Cochlear Nucleus. J Neurophysiol 89(6):3070–3082. https://doi.org/10.1152/jn.00125.2002

Rothman JS, & Manis PB (2003b) Kinetic Analyses of Three Distinct Potassium Conductances in Ventral Cochlear Nucleus Neurons. J Neurophysiol 89(6):3083–3096. https://doi.org/10.1152/jn.00126.2002

Rothman JS, & Manis PB (2003c) The Roles Potassium Currents Play in Regulating the Electrical Activity of Ventral Cochlear Nucleus Neurons. J Neurophysiol 89(6):3097–3113. https://doi.org/10.1152/jn.00127.2002

Rudnicki M, & Hemmert W (2017) High Entrainment Constrains Synaptic Depression Levels of an In vivo Globular Bushy Cell Model. Front Comput Neurosci 11(March):1–11. https://doi.org/10.3389/fncom.2017.00016

Scott LL, Hage TA, & Golding NL (2007) Weak action potential backpropagation is associated with high-frequency axonal firing capability in principal neurons of the gerbil medial superior olive. J Physiol 583(2):647–661. https://doi.org/10.1113/jphysiol.2007.136366

Scott LL, Mathews PJ, & Golding NL (2010) Perisomatic Voltage-Gated Sodium Channels Actively Maintain Linear Synaptic Integration in Principal Neurons of the Medial Superior Olive. J Neurosci 30(6):2039–2050. https://doi.org/10.1523/JNEUROSCI.2385-09.2010

Shinn-Cunningham BG, Zurek PM, Durlach NI, & Clifton RK (1995) Cross-frequency interactions in the precedence effect. J Acoust Soc Am 98(1):164–171. https://doi.org/10.1121/1.413752

Smith PH, Joris PX, & Yin TCT (1993) Projections of physiologically characterized spherical bushy cell axons from the cochlear nucleus of the cat: Evidence for delay lines to the medial superior olive. J Comp Neurol 331(2):245–260. https://doi.org/10.1002/cne.903310208

Srinivasan S, Laback B, Majdak P, & Arnolder C (2020) Improving Interaural Time Difference Sensitivity Using Short Inter-pulse Intervals with Amplitude-Modulated Pulse Trains in Bilateral Cochlear Implants. JARO - J Assoc Res Otolaryngol. https://doi.org/10.1007/s10162-020-00743-6

Srinivasan S, Laback B, Majdak P, & Delgutte B (2018) Introducing Short Interpulse Intervals in High-Rate Pulse Trains Enhances Binaural Timing Sensitivity in Electric Hearing. JARO - J Assoc Res Otolaryngol 19(3):301–315. https://doi.org/10.1007/s10162-018-0659-7

Stange A, Myoga MH, Lingner A, Ford MC, Alexandrova O, Felmy F, … Grothe B (2013) Adaptation in sound localization: from GABA B receptor-mediated synaptic modulation to perception. Nat Neurosci 16(12):1840–1847. https://doi.org/10.1038/nn.3548

Stecker GC (2020) Modeling “Straightness” Versus “Briefness:” Do Adapting Neural Models Account for Temporal Weighting and Bandwidth Effects on Binaural Sensitivity? ARO Midwinter Meeting, San Jose, CA, USA.

Stimberg M, Brette R, & Goodman DFM (2019) Brian 2, an intuitive and efficient neural simulator. ELife 8:1–41. https://doi.org/10.7554/eLife.47314

Traer J, & McDermott JH (2016) Statistics of natural reverberation enable perceptual separation of sound and space. Proc Natl Acad Sci 113(48):E7856–E7865. https://doi.org/10.1073/pnas.1612524113

Wallach H, Newman EB, & Rosenzweig MR (1949) A precedence effect in sound localization. Am J Psychol 62(3):315–336. https://doi.org/10.1121/1.1917119

Wang Y, & Manis PB (2008) Short-Term Synaptic Depression and Recovery at the Mature Mammalian Endbulb of Held Synapse in Mice. J Neurophysiol 100(3):1255–1264. https://doi.org/10.1152/jn.90715.2008

Xia J, Brughera A, Colburn HS, & Shinn-Cunningham B (2010) Physiological and psychophysical modeling of the precedence effect. JARO - J Assoc Res Otolaryngol 11(3):495–513. https://doi.org/10.1007/s10162-010-0212-9

Yang H, & Xu-Friedman MA (2009) Impact of Synaptic Depression on Spike Timing at the Endbulb of Held. J Neurophysiol 102(3):1699–1710. https://doi.org/10.1152/jn.00072.2009

Yang H, & Xu-Friedman MA (2015) Skipped-Stimulus Approach Reveals That Short-Term Plasticity Dominates Synaptic Strength during Ongoing Activity. J Neurosci 35(21):8297–8307. https://doi.org/10.1523/JNEUROSCI.4299-14.2015

Yin CT, & Chan JCM (1990) Interaural time sensitivity in medial superior olive of cat. J Neurophysiol 64(2):465–488. https://doi.org/10.1152/jn.1990.64.2.465

Zheng X, & Voigt HF (2006a) A modeling study of notch noise responses of type III units in the gerbil dorsal cochlear nucleus. Ann Biomed Eng 34(4):697–708. https://doi.org/10.1007/s10439-005-9073-5

Zheng X, & Voigt HF (2006b) Computational model of response maps in the dorsal cochlear nucleus. Biol Cybern 95(3):233–242. https://doi.org/10.1007/s00422-006-0081-9

Zhou Y, Carney LH, & Colburn HS (2005) A Model for Interaural Time Difference Sensitivity in the Medial Superior Olive: Interaction of Excitatory and Inhibitory Synaptic Inputs, Channel Dynamics, and Cellular Morphology. J Neurosci 25(12):3046–3058. https://doi.org/10.1016/j.physa.2016.12.040

Zilany MSA, Bruce IC, & Carney LH (2014) Updated parameters and expanded simulation options for a model of the auditory periphery. J Acoust Soc Am 135(1):283–286. https://doi.org/10.1121/1.4837815

Zilany MSA, Bruce IC, Nelson PC, & Carney LH (2009) A phenomenological model of the synapse between the inner hair cell and auditory nerve: Long-term adaptation with power-law dynamics. J Acoust Soc Am 126(5):2390–2412. https://doi.org/10.1121/1.3238250

Zilany MSA, & Carney LH (2010) Power-law dynamics in an auditory-nerve model can account for neural adaptation to sound-level statistics. J Neurosci 30(31):10380–10390. https://doi.org/10.1523/JNEUROSCI.0647-10.2010

